# Defining influenza-specific B cells in vaccine responders, non-responders and influenza breakthrough infections

**DOI:** 10.64898/2026.03.19.710321

**Authors:** Lukasz Kedzierski, Ashleigh I Holloway, Jennifer R Habel, Louise C Rowntree, Shihan Li, Hayley A McQuilten, Fiona James, Lydia Murdiyarso, Ilariya Tarasova, Nancy HL Leung, Yuyun Chen, Dennis KM Ip, Stephen J Kent, Malik Peiris, Steven Y C Tong, Allen C Cheng, Tom C Kotsimbos, Jason A Trubiano, Jan Schroeder, Adam K Wheatley, Benjamin J Cowling, Sophie A Valkenburg, Thi H O Nguyen, Katherine Kedzierska

**Author notes:** Authors contributed equally (Lukasz Kedzierski, Ashleigh I Holloway, Katherine Kedzierska). Authors contributed equally (Benjamin J Cowling, Sophie A Valkenburg, Thi H O Nguyen, Katherine Kedzierska).

## Abstract

Although seasonal influenza vaccination programs are effective at a population level, our data from inactivated influenza vaccine (IIV) cohorts in years 2015-2022 reveal that 50-60% of individuals do not seroconvert following immunization. The underlying mechanisms of vaccine non-responsiveness are far from understood. In this study, we sought to define key determinants of optimal B cell immune responses elicited by seasonal influenza vaccination, and to explore why some individuals fail to elicit humoral immunity following immunization. Immune responses associated with seroconversion and vaccine failure from individuals immunized with IIVs were compared at cellular and molecular levels using single-cell transcriptomics. We analyzed HA-specific B cell immunity across vaccine-responders, breakthrough infections and patients hospitalized with acute influenza. Droplet-based single-cell RNA sequencing and VDJ-sequencing of influenza-specific B cells from stored PBMCs was performed using 10x Genomics. Our results show that atypical B cells are the major subset of B cell responses in vaccine non-responders on day 28 post-vaccination. Conversely, individuals who seroconvert had diverse B cell phenotypes. The use of recombinant influenza-specific HA probes allowed us to dissect expression patterns on influenza HA-specific B cells. We found that HA-specific B cells of vaccine non-responders for A/H1N1 and A/H3N2 components displayed elevated atypical-like markers (CD11c, FcRL-5) at baseline, compared to responders. Analysis of differentially expressed genes (DEGs) between responders and non-responders identified differential expression of *HLA-DR*, *CD74*, *CD83*, and *CXCR3* genes. We subsequently demonstrated reduced frequencies of HLA-DR-, CD74- and CD83-expressing B cells in patients hospitalized with influenza, compared to healthy participants. Hospitalized influenza patients also had significantly higher proportions of atypical CD21^-^CD27^-^ B cells. Overall, our data demonstrate an association between elevated frequencies of atypical-like B cells with both lack of seroconversion following immunization and severe influenza infection. These findings broaden our understanding of humoral immunity in influenza vaccination and infection, providing novel insights for vaccination strategies and design.

## INTRODUCTION

Influenza A and B viruses (IAV and IBV) co-circulate annually, causing significant clinical and economic burden globally ^1^. Influenza vaccines, including inactivated influenza vaccines (IIVs), are currently the most effective public health intervention, thus key for inducing influenza-specific immunity against seasonal influenza viruses ^2^. The constant evolution of circulating influenza strains necessitates annual updates of influenza vaccine components across both the Northern and Southern hemispheres ^3^. Since 2012, most IIVs had been quadrivalent, containing four viral components: two IAVs (A/H1N1 and A/H3N2) and two IBVs (B/Victoria and B/Yamagata lineages) ^4^. During 2024-2025, the WHO advised that the inclusion of B/Yamagata strain in the influenza vaccine was no longer required due to the disappearance of B/Yamagata viruses during the COVID-19 pandemic ^5^.

Influenza vaccines reduce viral infections, transmission and disease severity ^6^. Vaccine efficacy is evaluated using vaccine effectiveness (VE), measuring how efficaciously vaccines reduce mortality and morbidity linked to influenza-related disease severity ^7^. VE varies from season to season, largely depending on the antigen match between vaccine and circulating strains ^8^. For well-matched vaccine formulations, it is estimated that the overall effectiveness of the influenza vaccine is between 40-42% ^9^. In case of a mismatch, VE can be as low as 10-20% ^10^. Efficacy also varies across vaccine components, with the lowest being for A/H3N2 strains ^11^, linked to a higher mutation rate of A/H3N2 viruses compared to A/H1N1 and IBVs ^12^, and mutations to the H3 component in egg-based viral passage ^13^. This was exemplified by the Australian 2017 influenza season, with the overall VE of IIV at 33%, mostly affected by a very low (10%) VE estimate for the H3 component ^14^. Pre-existing immunity, lack of seroconversion, comorbidities and age can all impact humoral responses to influenza vaccination ^15–18^.

Immune responses to IIVs involve both humoral and cellular immunity ^19^; especially B cells and T follicular helper cells ^20,21^. Antibody responses are predominantly measured using the haemagglutinin (HA) inhibition (HAI) assay, which detects HA-specific antibodies. HAI titers define correlates of protection against influenza, with a post-vaccination HAI titer of 40 correlating with 50% protection ^22^. Vaccine responders are defined by their 4-fold or more increase in HAI titers following IIV. Influenza-specific B cells generated by IIVs include memory B cells, class-switched memory B cells and antibody-secreting B cells. Recently, atypical memory B cells (atB cells), defined by their T-bet^+^CD11c^+^FcRL5^+^CD21^lo^ expression, were also reported following IIVs ^23^. AtB cells feature functionally-diverse subsets ^24^, and are also known for their responses in chronic infections and autoimmunity ^25^. In vaccination, there are conflicting data on the role of atB cells, with studies indicating that atB cells are predictors of long-lived antibody responses following IIV ^26^, or in contrast leads to diminished antibody responses after vaccination in older individuals ^27,28^.

In our study, we defined immunity in vaccine responders and non-responders following IIV in years 2015-2022, at cellular and molecular levels. Our data indicate that in Australia, 50-60% of participants do not experience 4-fold increase in HAI titers between day 0 (d0) and day 28 (d28) post-vaccination. A substantial proportion of those non-seroconverters (60%), however, had high HAI titers (>40) at baseline (d0), indicating pre-existing humoral immunity. We also analyzed influenza-specific immunity in participants with well-documented breakthrough infections following vaccination as well as in patients hospitalized with influenza. Our single cell RNA sequencing (scRNA-seq) on influenza HA-specific B cells and bulk B cells from vaccinated individuals at d0, d7 and d28 post-IIV, and individuals with breakthrough infections at baseline, post-vaccination and post-infection, revealed differential B cell immune features between IIV responders, non-responders and patients with breakthrough infections. Interestingly, atB cells were substantially higher in non-responders towards A/H1N1 and A/H3N2 at baseline, as well as in participants following breakthrough influenza virus infection. Moreover, we detected two separate clusters of HA-specific atB cells in longitudinal samples from individuals with breakthrough infections, one post-IIV, and the second following IAV infection. Our study also demonstrates, for the first time, that hospitalized patients with severe influenza display high levels of atB cells, a finding previously not described for acute influenza. This was associated with reduced CD74^+^ and HLA-DR^+^ expression on B cells in hospitalized patients, suggesting that B cells in severe influenza might have reduced activation and proliferation, leading to suboptimal immunity towards influenza viruses.

## RESULTS

### High proportion of non-responders in healthy adult IIV cohorts

We recruited 162 healthy adult participants vaccinated with IIV between 2015 and 2022. Trivalent IIV was used in 2015, whereas quadrivalent IIV was introduced in Australia from 2016 onwards (Fig. 1A). Median age of vaccinees was 34 years; 35.5 years for females and 31.5 years for males (Fig. 1B). We collected blood for PBMCs and sera at baseline, day 0 (d0), and d28 following vaccination to define influenza-specific B cells and antibodies.

**Figure 1.**
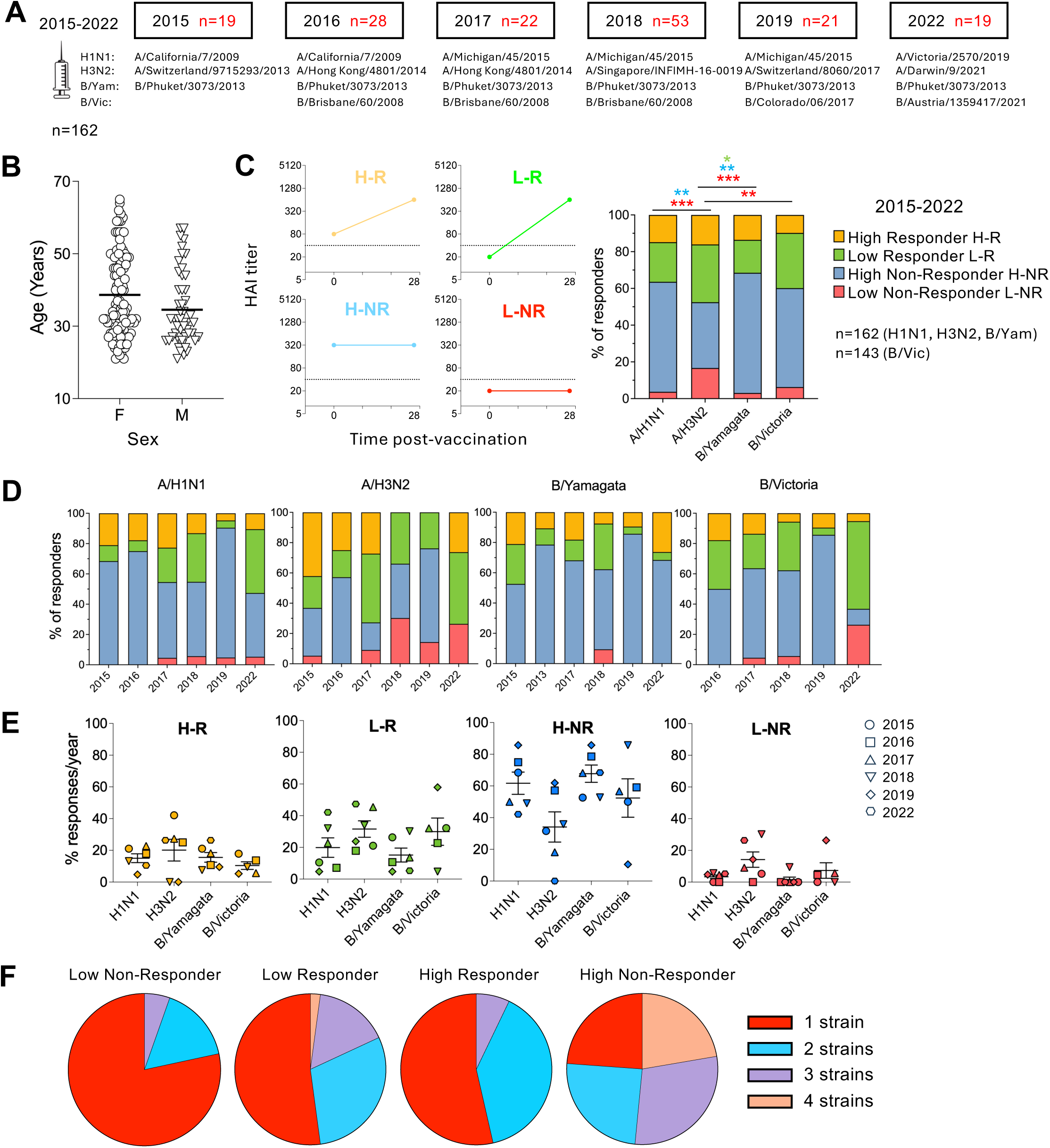
High proportion of Non-Responders in healthy adult IIV cohorts. (A) Vaccine composition and number of participants in healthy volunteer vaccination studies between 2015-2022. (B) Age and sex distribution of all participants. Line represents mean. C) HAI titer visualization used to stratify participants into High Responders (H-R), Low Responders (L-R), High Non-Responders (H-NR) and Low Non-Responders (L-NR) (left). Combined proportions of these four groups for each IIV component (right). Statistical significance was assessed by Chi square test (*P<0.05, **P<0.01, ***P<0.001). (D) Proportions of responders for each vaccine component across the years. (E) Proportions of different Responder/Non-Responder groups across each vaccine components per year. Line represents mean ± SEM. (F) Overall proportions of participants with different seroconversion status showing their responsiveness or non-responsiveness to vaccine components.

We first analyzed IIV participants for their vaccine responsiveness based on HAI titers detected on d28 following vaccination in comparison to d0 baseline titers. We stratified participants into four Responder and Non-Responder groups based on their HAI seroconversion status as well as pre-existing humoral immunity. Seroconverted Responders (R) were classified as individuals exhibiting a 4-fold or more increase in HAI titers following IIV between d0 and d28 ^22^. Further classification of Responders was based on whether their baseline HAI antibody titers were low (<40; Low Responders (L-R)) or high (≥40; High Responders (H-R) (Fig. 1C; left panel). Conversely, Non-Responders (NR) were classified as individuals lacking a 4-fold or more increase in HAI titers following IIV between d0 and d28. High Non-Responders (H-NR) had baseline HAI titers ≥40 and did not seroconvert, whereas Low Non-Responders (L-NR) had baseline HAI titers <40 and did not seroconvert.

Our initial analyses across vaccination years and IIV components found that a mean of 61.2% vaccinees did not respond to IIV based on HAI seroconversion data (H-NR and L-NR grouped together; Fig. 1C). Considering the IIV component, 63.6% vaccinees did not respond to A/H1N1, 52.5% to H3N2, 68.5% to B/Yamagata and 60.1% to B/Victoria. Given such a high proportion of vaccinees not seroconverting to IIV, we further defined IIV participants according to their HAI antibody baseline levels, as outlined above. We found the highest proportion (16.67%) of Low Non-Responders to A/H3N2 antigen; significantly higher when compared to the other vaccine components (A/H1N1=3.7%, B/Yamagata=3.09%, B/Victoria=6.29%; P<0.005) (Fig. 1C; right panel). Consequently, we observed the lowest proportion of High Non-Responder participants (35.8%; no seroconversion but with baseline antibody levels) to A/H3N2 vaccine component; significantly lower than to the A/H1N1 component (P=0.0097, 59.9%) and B/Yamagata component (P=0.0021, 65.4%), but no different to B/Victoria (53.9%).

With respect to Responder groups, the only difference was observed for the Low-Responder group where A/H3N2 (31.5%) was higher than B/Yamagata components (17.9%) (P=0.0274). No differences were found in proportions of High-Responder participants between all four IIV components, ranging between 9.8% to 16.1% (Fig. 1C; right panel).

Further analyses across individual IIV components and across different years revealed similar frequencies of our four defined Responder and Non-Responder groups. High-Non-Responder participants accounted for the largest proportion of all groups, although there was a consistent presence of a Low Non-Responder group in the A/H3N2 analysis with each year, except for 2016 (Fig. 1D). None of the differences, however, reached statistical significance when comparisons between proportion of each group per vaccination year were assessed for each vaccine component (Fig. 1E). Low Non-Responders predominantly showed lack of seroconversion to a single influenza strain included in IIV, whereas High Non-Responders lacked responsiveness at almost equal proportions of individuals not seroconverting to either 1, 2, 3 or all vaccine components (Fig. 1F).

These data suggest that pre-existing immunity acquired by previous influenza vaccination or infection, defined here by baseline HAI antibody levels, plays an important role in seroconversion following IIV in healthy individuals.

### Single-cell RNA-seq in IIV-immunized participants reveal memory and atypical B cell clusters

To gain in-depth insights into B cell responses in the Non-Responder and Responder IIV groups, we performed single-cell RNA sequencing (RNA-seq) using recombinant hemagglutinin (rHA) probes directed towards all four IIV vaccine components ^21,29,30^. We selected 6 participants from the 2022 cohort for analyses of their HA-specific B cells across all four IIV components (thus 24 samples of HA-specific B cell populations), based on their diverse responder classification (Fig. 2A). We focused on those 24 HA-specific B cell populations in IIV-vaccinated individuals across 3 timepoints: d0, d7 and d28 post-vaccination. rHA-specific B cells from each participant and timepoint were isolated for single-cell RNAseq from PBMCs using surface marker antibodies, barcoded rHA probes and barcoded cellular indexing of transcriptomes and epitopes (CITE-Seq) antibodies. RNA and protein expression data were used to generate a weighted nearest neighbour uniform manifold approximation projection (wnnUMAP) of B cells. Following the quality control, nine unsupervised B cell clusters were generated (Fig. 2B). Clusters were manually annotated using previously published gene signatures (Fig. 2C) ^23,31,32^. We then used 9 genes (*ITGAX, FCRL5, IGHD, CD27, CXCR4, CXCR3, TCL1A, EBI3, IGHG1*) and 5 surface proteins (CD11c, CD21, CD27,CXCR3, CXCR5) expression profiles to further define the B cell subset in each clusters (Fig. 2D, E). B cells were assigned as naïve (B naïve; clusters 0, 4, 6), memory (B mem; clusters 1, 2, 3, 5), plasmablasts (PB; cluster 8) and atypical memory (atB; cluster 7). On average, each participant contributed similarly to B cell clusters, except for Participant 4 having the highest proportion of cells in cluster 7 (atB), whereas cluster 0 (B naïve) was dominated by Participants 1, 2 and 3 (Fig. 2F).

**Figure 2.**
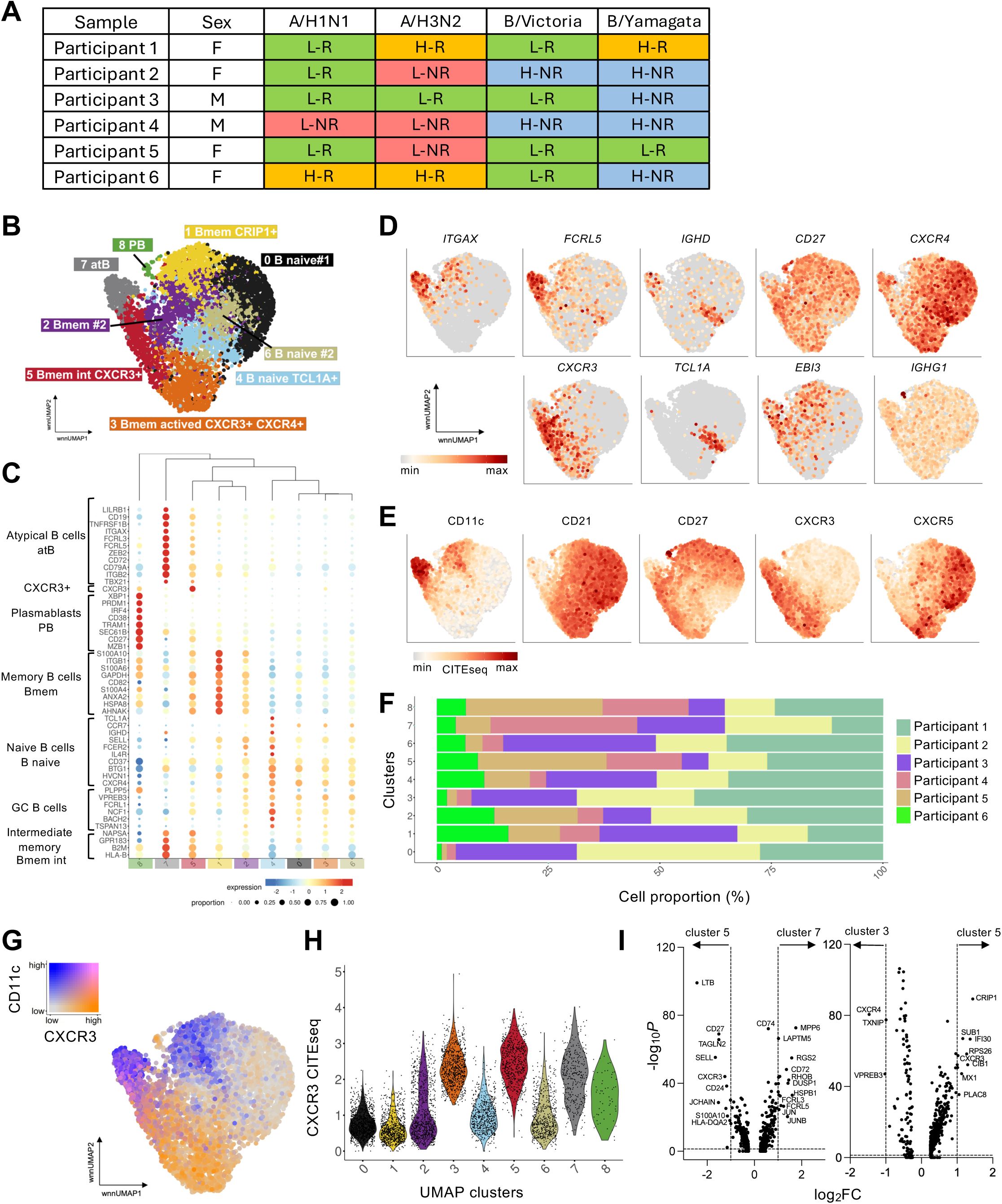
scRNA-seq transcriptional profiling of healthy vaccinated participants. (A) IIV participants used in scRNA-seq analysis. rHA^+^ B cells across all 4 vaccine components were sorted for 6 participants; leading to 24 rHA^+^ B cell samples. (B) wnnUMAP depicting 9 major clusters of bulk B cells with annotations shown. (C) Hierarchically clustered bubble plot of selected marker gene signatures for each cluster. Colour depicts mean gene expression and dot size depicts frequency of cells expressing a gene in a cluster. (D-E) Feature plots of (D) GEX and (E) CITEseq showing distinguishing B cell markers in the total B cell dataset. (F) Proportion of cells in each wnnUMAP cluster per participant. (G) wnnUMAP depicting CD11c and CXCR3 protein co-expression across the clusters. (H) Violin plots showing CXCR3 protein expression per cluster. (I) Volcano plots of differential gene expression between cluster 5 and 7 (left) and clusters 3 and 5 (right). Horizontal dashed lines depict statistical significance threshold, vertical dashed lines indicate the log_2_FC cutoff.

Further in-depth analysis focused on differences between B mem clusters 3 and 5 and atB cluster 7, which showed high expression levels of CXCR3 at the gene and protein levels (Fig. 2D, E). Co-expression of CD11c and CXCR3 is thought to mark recently activated atypical B cells ^23^. Cells in atB cluster 7 were predominantly CD11c single-positive, with a large proportion of cells being CD11c and CXCR3 double-positive (Fig. 2G, H). This, in addition to being CD21^-^CD27^int^, and expression of *ITGAX* and *FCRL5*, indicated that cluster 7 represents “classical” atB cells (Fig. 2D,E). In contrast, B mem clusters 3 and 5 showed higher CXCR3 expression levels but low CD11c expression (Fig. 2D,E). Differentially expressed gene (DEG) analysis further showed that *CXCR3* and *CD27* were highly upregulated in cluster 5, whereas *FCRL3, FCRL5* and *CD72* were highly expressed in cluster 7 (Fig. 2I left panel). Comparison of DEGs between cluster 3 and 5, positive for CXCR3, indicated that *CXCR4* was the most significantly upregulated gene in cluster 3, whereas cluster 5 featured *CXCR3* and *CRIP1* expression (Fig. 2I right panel). *CRIP1* was also highly expressed in cluster 1 (memory cells), previously shown to be upregulated in memory B cells ^33,34^. Additionally, cluster 5 showed high expression of the chemokine receptor, CXCR5 (Fig. 2D).

### Differential distribution of influenza HA-specific B cells between vaccine Responders and Non-Responders

We sought to next define transcriptomic signatures of HA-specific B cells in Responders and Non-Responders following IIV immunization. We used barcoded APC-labelled rHA probes to A/Sydney/5/2021 H1N1 (H1), A/Darwin/9/2021 H3N2 (H3), B/Austria/1359417 (B/Vic) and B/Phuket/3073/2013 (B/Yam) to identify influenza-specific rHA^+^ B cells towards all four IIV components at different timepoints post-IIV. rHA^+^CD71^+^IgD^-^CD19^+^ B cells towards combined four rHA probes were readily detected in all participants at baseline, indicating past infections or vaccinations (Fig. 3A, B). Following IIV immunization, rHA-specific B cell population across all 4 rHA probes expanded in 50% of participants by d28, remained unchanged in one and contracted in two remaining participants (Fig. 3B), illustrating the spectrum of responses to IIV.

**Figure 3.**
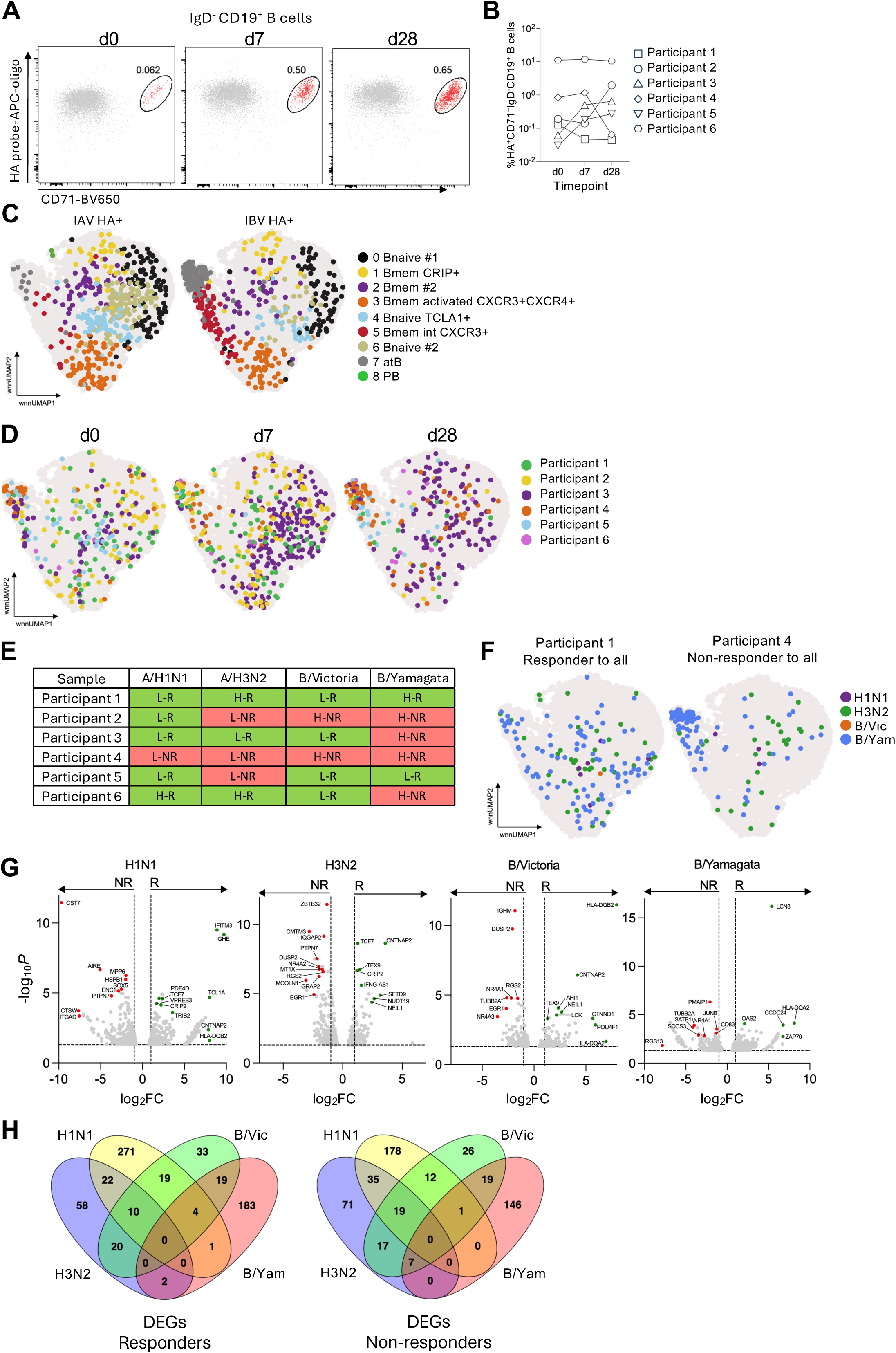
Differential distribution of rHA-specific B cells between vaccine Responders and Non-Responders. (A) Representative FACS plots showing total rHA^+^CD71^+^ activated B cells (Participant 3). rHA^+^ cells of all 4 specificities were labelled with different APC-oligo barcodes, cells were gated on IgD^-^CD19^+^ B cells. (B) Frequencies of total rHA^+^CD71^+^IgD^-^CD19^+^ B cells per participant across timepoints. (C) wnnUMAP depicting global distribution of IAV^+^ and IAV^+^ rHA^+^ B cells across 9 clusters. Colours denote different clusters. (D) wnnUMAP showing rHA^+^ B cells at different time points post-vaccination. Colours depict individual participants. (E) Table showing 6 participants used in scRNA-seq experiment stratified as either Responders (R, green) or Non-Responders (NR, red) regardless of their initial HAI titer. (F) wnnUMAP showing rHA^+^ B cells distribution for Participant 1 (seroconverted to all IIV components, left) and Participant 4 (no seroconversion to all IIV components, right). Colour depicts influenza strain specificity. (G) Volcano plots of differential gene expression between Responders (R) and Non-Responders (NR) for each IIV vaccine component. Only genes with significant differential expression and ≥2-fold change in expression were plotted. Horizontal dashed lines depict statistical significance threshold, vertical dashed lines indicate the log_2_FC cutoff. Coloured dots depict genes of interest in R (green) or NR (red). (H) Venn diagrams showing the number of overlapping genes identified by scRNA-seq in Responders (left) and Non-Responders (right) between different strains specificities.

IAV- and IBV-specific B cells displayed different distribution across clusters, particularly evident in cluster 5 (Bmem intermediate CXCR3+) and 7 (atB), mostly populated by IBV+ B cells (Fig. 3C). Conversely, cluster 4 (B naïve TCLA1+) and cluster 6 (B naïve #2) were enriched in IAV+ B cells. We subsequently analyzed all rHA+ B cells across 3 timepoints as well as individual participant’s contribution to B cell clusters (Fig. 3D). Throughout the time-course, a changing pattern of distribution was observed between the participants, with evident expansion of influenza-specific B cells on d7 post-IIV. Notably, cluster 7 (atB cells) was populated by B cells derived from different participants across all timepoints, with the largest contribution from Participant 4 on d0 and d28 (Fig. 3D). This indicates that atypical B cells are an integral component of B cell response to IIV immunization, as previously reported ^23^.

To identify differential transcriptomic signatures of rHA+ B cells following IIV immunisation in Responders and Non-Responders, we grouped participants into responders (R; in green) and Non-Responders (NR; in red) regardless of their initial HAI status (Fig. 3E), then performed a pseudo-bulk differential gene expression analysis between Responder and Non-Responder groups for each IIV component. Notably, this grouping included Participant 1, who responded to all IIV components and Participant 4, who was Non-Responder to all four IIV components. The remaining four participants displayed combinations of baseline HAI titers and seroconversion status. Interestingly, Participant 1 showed a broad distribution of total rHA+ B cells (all timepoints combined) throughout clusters, whereas Participant 4 mainly contributed to cluster 7 (Fig. 3F). The rHA+ cells were mostly of H3 or B/Yam specificity.

We identified a total of 634 significant DEGs in H1N1 group, with 355 genes upregulated in H1 Responder group (327 genes with >2-fold increase) and 279 genes upregulated in Non-Responders (245 genes with >2-fold increase) (Fig. 3G; Supplementary Table S1). The most significantly upregulated genes in the H1 Responders included *IFITM3*, *IGHE*, *TCL1A*, *HLA-DQB2* and *CNTNAP2*, whereas Non-Responders upregulated *CST7*, *AIRE*, *MPP6*, *HSPB1* and *SOX5.* In the H3-specific comparison, we identified a total of 491 significant DEGs with 190 genes upregulated in H3 Responders (112 genes with >2-fold increase) and 301 genes upregulated in Non-Responders (149 genes with >2-fold increase) (Fig. 3G; Supplementary Table S2). Some of the most significantly upregulated genes in Responders included *CNTNAP2*, *TCF7*, *TEX9*, *CRIP2* and *IFNG-AS1*, whereas Non-Responders significantly upregulated *ZBTB32*, *CMTM3*, *PTPN7*, *DUSP2* and *NR4A2*.

Similar analyses for B/Vic specificity resulted in 299 significant DEGs with 140 genes upregulated in Responders (105 genes with >2-fold increase, including *HLA-DQB2, CNTNAP2, AHI1, TEX9, NEIL1* or *LCK*) and 159 genes upregulated in Non-Responders (101 genes with >2-fold increase featuring *IGHM, DUSP2, NR4A1, NR4A3, TUBB2A*) (Fig. 3G; Supplementary Table S3). For B/Yamagata-specific B cell, we detected 474 significant DEGs with 254 upregulated in Responders (209 with >2-fold increase such as *LCN8, HLA-DQA2, OAS2, ZAP70*) and 220 in Non-Responders (173 genes with >2-fold increase including *PMAIP1, C5, TUBB2A, SATB1, NR4A1, CD83*) (Fig. 3G; Supplementary Table S4).

We then looked for common gene signatures in response to the four vaccine components and compared all DEGs significantly upregulated in Responder groups (Fig. 3H, Supplementary Table S5). Although there were no common elements across all 4 IIV specificities, four genes were common in response to H1N1, B/Victoria and B/Yamagata; *AK8, LCK, SH2D3A, PRIM1*. A larger group of 10 genes were common in response to H1N1, H3N2 and B/Victoria and included *CNTNAP2, SETD9, NEIL1, XXYLT1, AC253572.2, IFNG-AS1, TSGA10, FGD6, EPHA4, ABHD5*. In Non-Responder group again, there was no common gene upregulation between all 4 IIV specificities, but 7 genes were common in response to H3N2, B/Victoria and B/Yamagata *TOP1MT, ATP13A3, SERTAD3, NR4A1, DUSP5, AREG, TUBB2A*), one gene in response to H1N1, B/Victoria and B/Yamagata (*RTP5*) and 19 genes were common in response to H3N2, H1N1 and B/Victoria (*TIMP1, CLEC2B, METRNL, LINC01128, KANSL1-AS1, CEMIP2, DGKG, RGS2, NR4A2, SERTAD1, TRIB1, AC104024.1, MT1X, GRAP2, PTPN7, SEMA4A, EGR1, CMTM3, MCOLN1*) (Fig. 3H, Supplementary Table S6).

### V(D)J gene usage and clonal expansion in influenza HA-specific B cells after IIV vaccination

To understand whether B cell receptor (BCR) clonality impacted the IIV Responder versus Non-Responder status, we analysed the V and J gene segments of BCRs, components of the heavy chain (IGH) playing an important function in shaping BCR repertoires. In particular, V gene segments can be associated with the HA protein ^35^. Analyses of A/H3N2-and B/Yamagata-specific BCR repertoires were performed in-depth (Fig. 4A), A/H1N1- and B/Victoria-specific BCR sequences were too low to perform meaningful BCR analyses. We found that A/H3N2-specific B cells of Responders had significantly increased the frequency of the heavy chain segment *IGHV3-23* (P=0.0325, Fisher’s exact test, unadjusted p-value) post-IIV immunization from d7 to d28 (Fig. 4B). Additionally, we found a trend for increased frequency of this segment between d7 and d28, compared to Non-Responders, albeit not statistically significant. This was not observed in B/Yamagata-specific B cells (Fig. 4B). Conversely, the A/H3N2-specific B cells of Non-Responders had significantly increased frequency of the light chain segment *IGKV3-20* (P=0.0327, Fisher’s exact test, unadjusted p-value) and *IGLV3-1* (P=0.0159, Fisher’s exact test, unadjusted p-value), which was not observed in the A/H3N2 Responders, nor the B/Yamagata-specific B cells. The differences in frequencies of *IGKV3-20* were also significantly different (P = 0.0002, Fisher’s exact test, unadjusted p-value) on d28 between Non-Responder and Responder groups (Fig. 4B).

**Figure 4.**
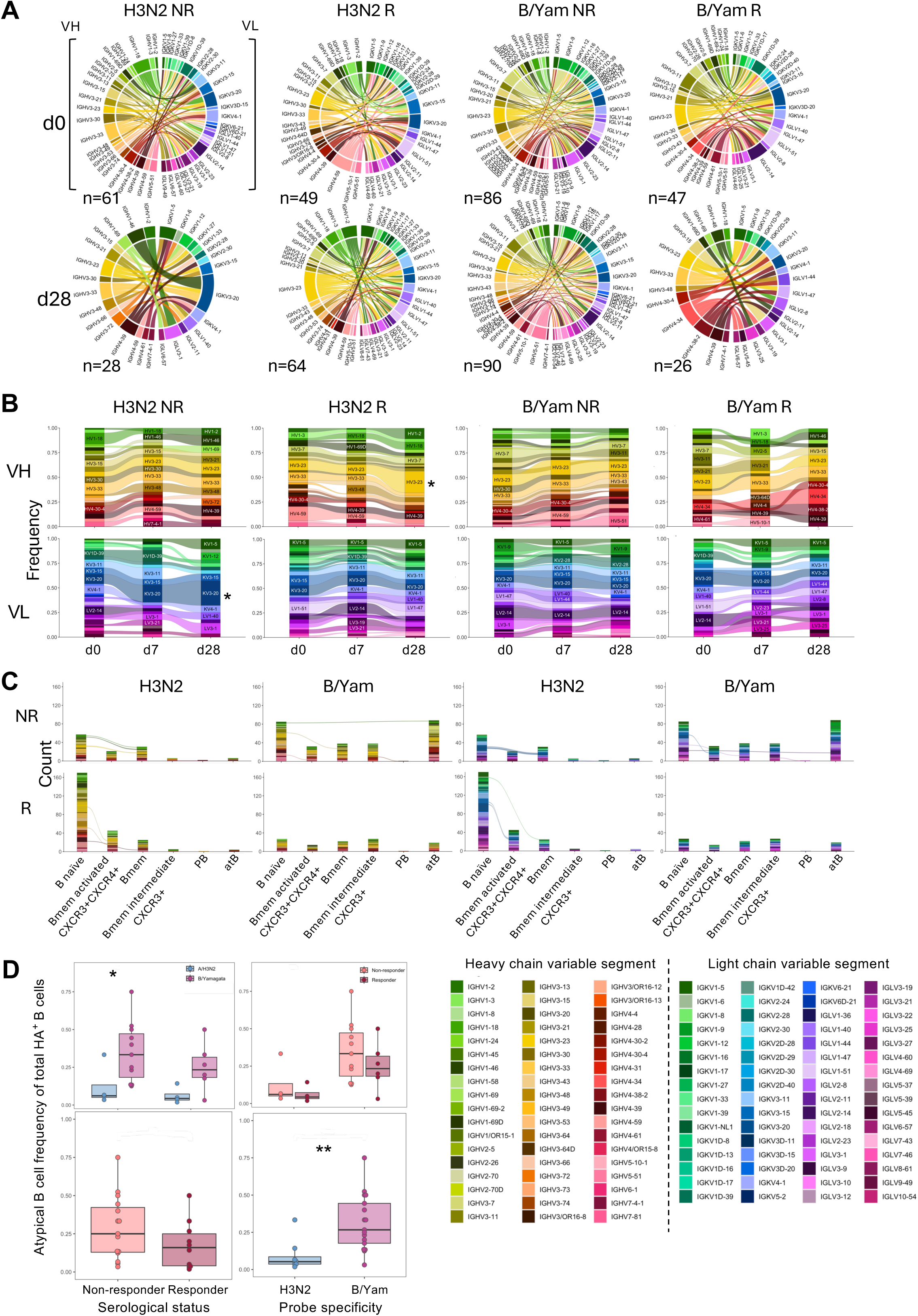
V(D)J analysis of BCR repertoires in IIV vaccinees. (A) Circos plots showing BCR heavy (VH) and light (VL) chain pairings for A/H3N2 and B/Yamagata specificities in Responders (R) and Non-Responders (NR). (B) Frequencies of VH and VL chains at d0, d7 and d28 post-vaccination for A/H3N2 and B/Yamagata specificities in NR and R groups. (C) VH (left) and VL (right) chain usage in A/H3N2 and B/Yamagata NR and R groups in B cell clusters identified in scRNA-seq. (D) Comparison of frequency of rHA-specific atypical B cells between NR and R for A/H3N2 and B/Yamagata specificities. Tukey box and whisker plots, boxes show 1^st^ to 3^rd^ quartile, line indicates median and whiskers extend to the maximum and minimum values in the data. Statistical significance was assessed using unadjusted Mann-Whitney test (*P<0.05, **P<0.01).

To define any differences in BCR usage across B cell clusters, we plotted the frequency of heavy (Fig. 4C left side) and light chain (Fig. 4C right side) variable segments for A/H3N2-specific and B/Yamagata-specific B cell populations for both Responders and Non-Responders. Highly diverse variable gene segment usage among each of the clusters was evident, but notably a significantly higher frequency of B/Yamagata-specific atypical B cells than A/H3N2 was detected (P=0.00468) (Fig. 4D). This was consistent with findings from the distribution of IAV- and IBV-specific B cells across clusters (Fig. 3C). Furthermore, the frequency of atypical B cells was higher, albeit not significantly, in Non-Responders compared to Responders for B/Yamagata, and the frequency of atypical B cells in B/Yamagata Non-Responders was significantly (P=0.0365) higher than A/H3N2 Non-Responders (Fig. 4D).

Overall, V(D)J analysis of BCR repertoires demonstrated that A/H3N2-specific B cell responses were highly polyclonal with a higher usage of the *IGHV2-23* and *IGKV3-20* segments in Responders and Non-Responders, respectively. These differences were not observed in responses to B/Yamagata strain, but IBV responses were dominated by atB cells in Non-Responders.

### Single-cell RNA-seq of B cells in participants with breakthrough influenza shows unique atypical B cell clusters

Having defined transcriptomic features of influenza rHA-specific B cells in vaccine Responders and Non-Responders, we further sought to understand transcriptomic signatures in vaccinated individuals who had breakthrough influenza virus infection within a year post-IIV immunization. We thus performed single-cell RNAseq on rHA-specific B cells in IIV-vaccinated individuals with a documented subsequent breakthrough infection, using rHA probes to IIV vaccine components (from a season prior to the breakthrough infection). We selected four participants enrolled in the RETAIN cohort, a unique multi-year longitudinal vaccination study to assess the efficacy of vaccination regimens in Hong Kong, an area with biannual influenza season. The longitudinal nature of this study allowed for the tracking of immune responses over time, including HAI titers, as well as detect influenza virus infection following immunization. We had access to three participants with breakthrough A/H3N2 infection and one participant with A/H1N1 infection (Fig. 5A). For 2 participants with A/H3N2 breakthrough infections, the HAI titer for the influenza virus infection strain, A/Singapore/2016 (H3N2), was greater than the 50% protective titer of 40. For the third H3N2 breakthrough participant as well as the A/H1N1 breakthrough infection participants (A/Brisbane/2018), the HAI titers were below 40 (Fig. 5A). 10x scRNA-seq analysis was performed on PBMCs from baseline (BL), post-vaccination (postvax) and post-infection timepoints.

**Figure 5.**
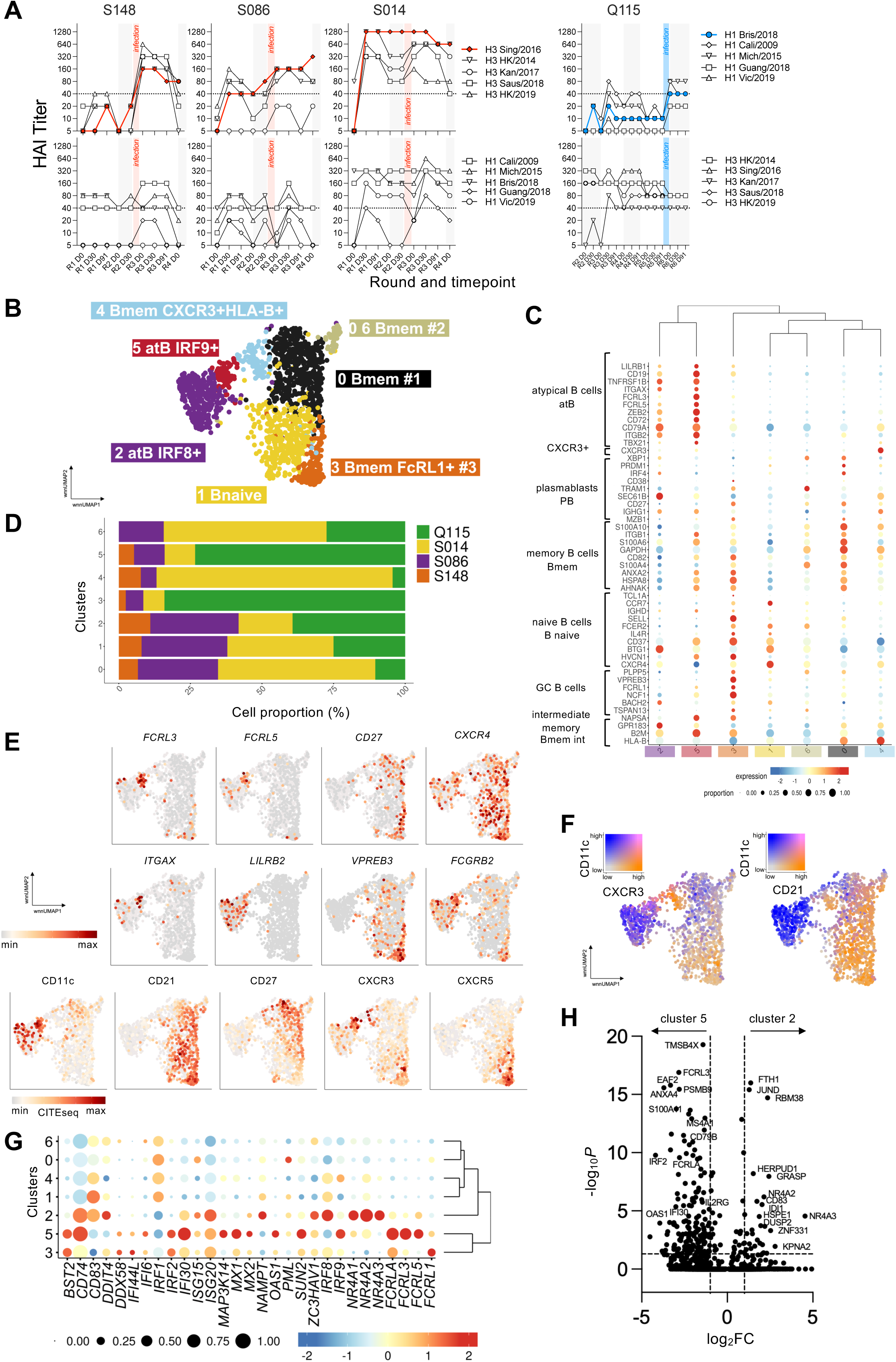
scRNA-seq transcriptional profiling of vaccinated participants with breakthrough infection. (A) Four vaccinees with a well-documented influenza infection history were used in our scRNA-seq. HAI titers to different strains of H1N1 and H3N2 influenza viruses across time prior to and post the infection are shown. Participants selected for scRNA-seq had a confirmed H3N2 infection (red) or H1N1 infection (blue). HAI titers to a breakthrough strain are shown as solid-coloured symbols. (B) wnnUMAP depicting 7 major clusters of bulk B cells with annotations shown. (C) Hierarchically clustered bubble plot of selected marker gene signatures for each cluster. Colour depicts mean gene expression and dot size depicts frequency of cells expressing a gene in a cluster. (D) The proportion of cells in each wnnUMAP cluster per participant. (E) Feature plots of GEX (upper 2 rows) and CITEseq (bottom row) showing distinguishing B cell markers in the total B cell dataset. (F) wnnUMAP depicting CD11c and CXCR3 and CD11c and CD21 protein co-expression across the clusters. (G) Hierarchically clustered bubble plot of selected gene signatures used to define clusters 2 and 5. Colour depicts mean gene expression and dot size depicts frequency of cells expressing a gene in a cluster. (H) Volcano plots of differential gene expression between cluster 2 and 5. Horizontal dashed lines depict statistical significance threshold, vertical dashed lines indicate the log_2_FC cutoff.

We identified seven B cells clusters based on gene expression signatures (Fig. 5B). Clusters 2 and 5 showed a gene expression pattern characteristic of atB cells (Fig. 5C) and, interestingly, participant Q115 with H1N1 breakthrough infection was the major contributor to cluster 5 and prominent in cluster 2 (Fig. 5D). Further analysis using 8 genes *(FCRL3, FCRL5, CD27, CXCR4, ITGAX, LILRB2, VPREB5, FCGRB2*) and 5 surface protein (CD11c, CD21, CD27, CXCR3, CXCR5) expression profiles in cluster 2 and 5 confirmed that both clusters were CD21- and CD27-double-negative and CD11c-positive (Fig. 5E, F). Interestingly, cluster 5 showed stronger *FCRL3* and *FCRL5* gene expression and higher levels of CXCR3. To further define these clusters, we analyzed expression pattern of a selection of ISGs, transcription factors and cellular receptors (Fig. 5G) and performed DEG analyses (Fig. 5H). Cluster 2 was defined as atB IRF8^+^, whereas cluster 5 was atB IRF9^+^. Moreover, cluster 2 was characterised by strong expression of *CD83* and members of nuclear receptor 4 (NR4) gene family (*NR4A1, NR4A2, NR4A3*). Conversely, cluster 5 was characterised by expression of genes encoding Fc receptor like proteins (*FCRLA, FCRL3, FCRL5*) and interferon-induced genes including *IRF2, IFI30, MX1* and *OAS1*. It is important to note that although atypical B cells following influenza vaccination have been previously reported ^20^, our study is first to describe atypical B cells in influenza virus infection.

Since Q115 was a major contributor to atB cells overall from clusters 2 and 5 combined (Fig. 5D), we analysed distribution of rHA+ B cells in this participant. Q115 demonstrated a robust response post-vaccination when compared to baseline rHA+ B cells, and no further rHA+ B cell expansion following breakthrough infection (Fig. 6A), with the overall wide distribution of B cells across the clusters (Fig. 6B). Interestingly, the majority of rHA+ B cells were of H3 specificity followed by H1 specificity, which post-vaccination were predominantly located in atB IRF8^+^ cluster 2, but then shifting to atB IRF9^+^ cluster 5 post-infection (Fig. 6C). We detected small expansion of H1/Victoria-specific rHA+ cells following vaccination within B mem #1 cluster 0, but no further expansion following the breakthrough infection. No expansion of the vaccine H1/Michigan specificity was observed following vaccination or breakthrough infection (Fig. 6C). Overall, this Q115 breakthrough infection case exemplifies differential transcriptomic features between rHA-specific atypical B cells following influenza vaccination and infection.

**Figure 6.**
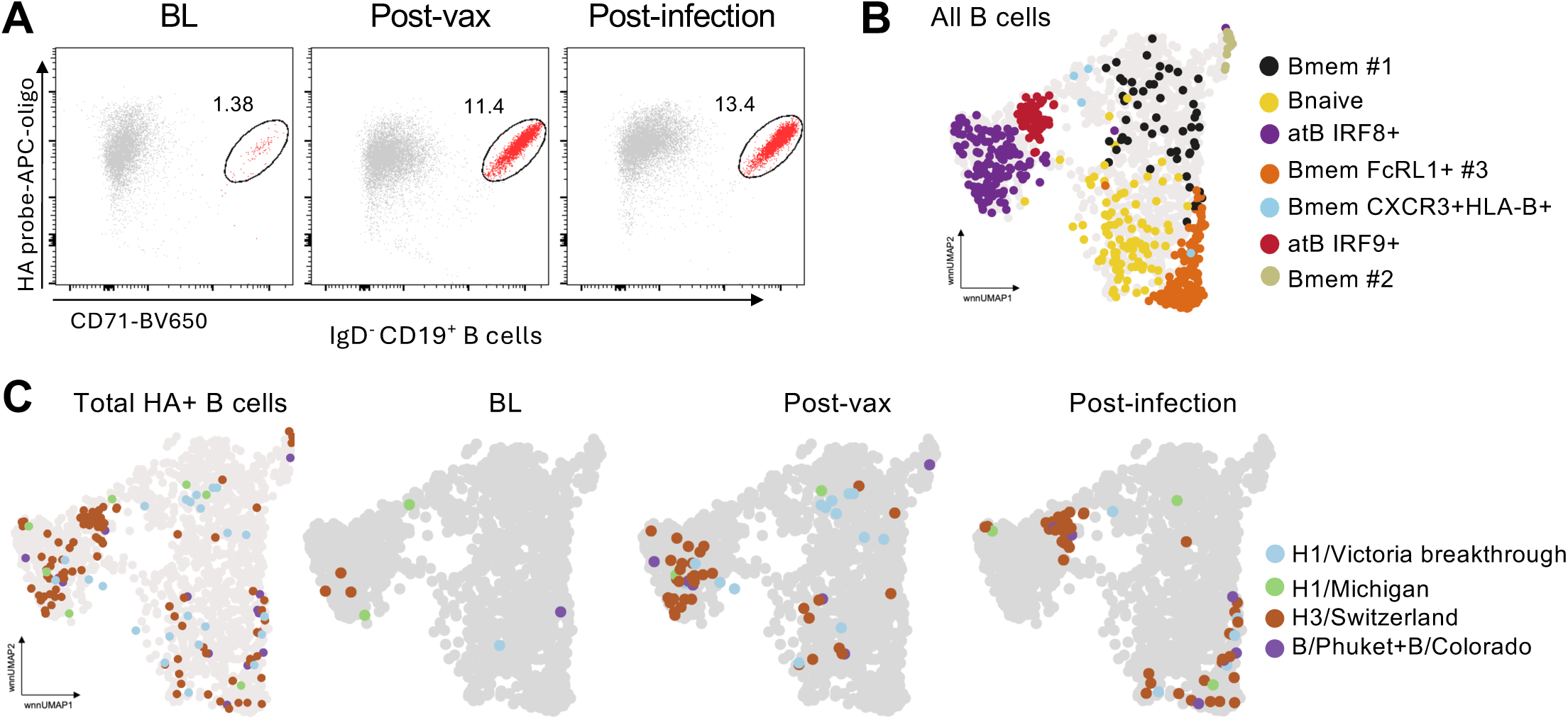
scRNA-seq transcriptional profiling of case study participant Q115 from RETAIN breakthrough infection cohort. (A) FACS plots of rHA probe staining at baseline (BL), post-vaccination (post-vax) and post-infection on live rHA^+^IgD^-^CD19^+^ B cell populations for IAV and IBV specificities pooled together on rHA-APC probes in participant Q115. (B) wnnUMAP depicting global distribution of pan B cells across 7 clusters. Colours denote different clusters. (C) wnnUMAP showing rHA^+^ B cells at different time points post-vaccination. Colours depict individual specificities.

### Influenza rHA-specific B cells in IIV Non-Responders display a predominant atypical B cell phenotype by flow cytometry

To verify our transcriptomics data at the protein level, we analysed influenza rHA-specific B cells by flow cytometry in 18 healthy vaccinated participants from the 2022 cohort (6 participants previously selected for scRNA-seq experiment (Fig. 2A) and 12 additional participants; Supplementary Table S7). All participants were stratified into Responder and Non-Responder groups based on their HAI titers. Influenza rHA-specific B cells were analysed at d0 and d28 post-vaccination (Fig. 7A). Analysis of rHA+ B cells across all specificities (H1, H3 and B) revealed a significantly increased frequency of rHA^+^IgD^-^ B cells following IIV in the Responder group (d0 vs d28; P=0.005), whereas rHA^+^IgD^-^ B cells remained unchanged in the Non-Responder group (Fig. 7B). Stratifying rHA^+^IgD^-^ B cells according to strain-specific responses, we found a significant increase in A/H3N2-specific B cells post-vaccination in Responders (d0 vs d28; P=0.0215), but not in Non-Responders. The same trend was observed in Responders for A/H1N1 and IBV specificities, albeit not statistically significant (Fig. 7B).

**Figure 7.**
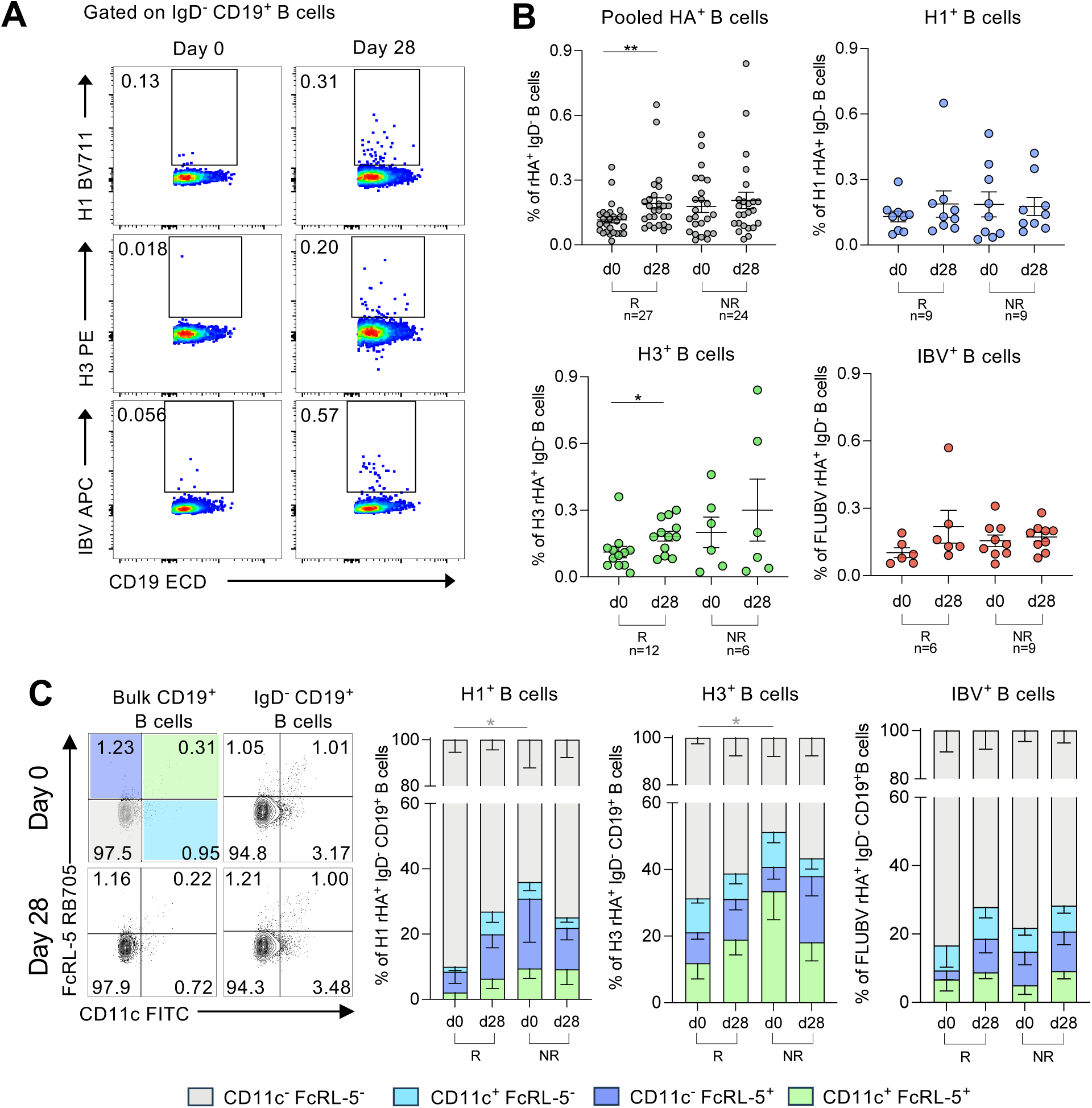
Influenza-specific B cell responses in healthy vaccinated individuals. (A) Representative FACS plots of rHA probe staining at d0 and d28 post-vaccination on live rHA^+^IgD^-^CD19^+^ B cell populations for IAV and IBV specificities (B/Yamagata and B/Victoria specificities were pooled together on a single rHA-APC probe). (B) Frequency of all influenza-specific (pooled probes) and H1-, H3- and IBV-specific rHA^+^IgD^-^CD19^+^ B cells at d0 and 28 days post-IIV vaccination in Responder (R) and Non-Responders (NR) for the respective vaccine component. As both B/Yamagata and B/Victoria rHA probes were conjugated to the same fluorochrome, IBV-specific rHA^+^IgD^-^ B cells were only analyzed for participants with the same Responder/Non-Responder status for both B cell lineages. Statistical significance was assessed using Mann-Whitney test (*P<0.05, **P<0.01). Bars indicate mean ± SEM. (C) Representative FACS plots of CD11c and FcRL-5 staining showing distribution of CD11c and FcRL-5 expression on bulk CD19^+^ and class-switched (IgD^-^CD19^+^) B cells (left). Frequency distribution of CD11c and FcRL-5 expression in influenza rHA-specific B cells within R and NR groups for respective vaccine components at d0 and d28 (right). Statistical significance was assessed using Mann-Whitney test for unpaired comparisons between groups (*P<0.05). Bars indicate mean ± SEM.

To further investigate the prevalence of atypical B cells in IIV participants, we phenotyped the rHA-specific B cells and investigated the distribution of atypical markers CD11c and FcRL-5 at d0 and d28 post-vaccination. CD11c and FcRL-5 gating was used to determine distribution of rHA^+^IgD^-^CD19^+^ B cells in Responders and Non-Responders, with double-positive CD11c^+^FcRL-5^+^ or single-positive, either CD11c^-^FcRL-5^+^ or CD11c^+^FcRL-5^-^, depicting atypical B cells (Fig. 7C). We observed higher overall expression of atB cell markers on IAV-specific B cells, exemplified by lower frequencies of CD11c^-^FcRL-5^-^ rHA-specific B cells. We detected significantly lower proportions of double-negative A/H1N1-(P=0.0396) and A/H3N2-specific (P=0.0167) non-atypical B cells in Non-Responders at d0, and thus higher total proportions of CD11c and/or FcRL-5-expressing atypical B cells, compared to Responders. No differences were found in IBV-specific B cells (Fig. 7C) or across the groups at d28 post-vaccination.

### Expression patterns of CD74, HLA-DR, CD83 and CXCR3 on rHA-specific B cells differ in Responders and Non-Responders

Analysis of scRNA-seq data form IIV (Fig. 2) and breakthrough infection cohorts (Fig. 5) revealed key B cell gene signatures within rHA-specific B cells, including CD74, CD83, HLA-DR and CXCR3. From the breakthrough infection cohort, CD74 was strongly expressed in clusters 2 and 5 (Fig. 5G) and CD83 was differentially expressed between atB clusters 2 and 5 (Fig. 5G and H). Whereas CXCR3 was expressed in B mem clusters 3 and 5, and atB cluster 7 from the vaccination cohort (Fig. 2G, H), and was previously reported to be associated with atB cells ^23,24,36^. Furthermore, HLA-DR expression on B cells was previously associated with influenza seroconversion ^37^. Thus, we selected these 4 markers for further phenotypic investigation of rHA-specific B cells in the context of IIV immunisation at the protein level (Fig. 8A).

**Figure 8.**
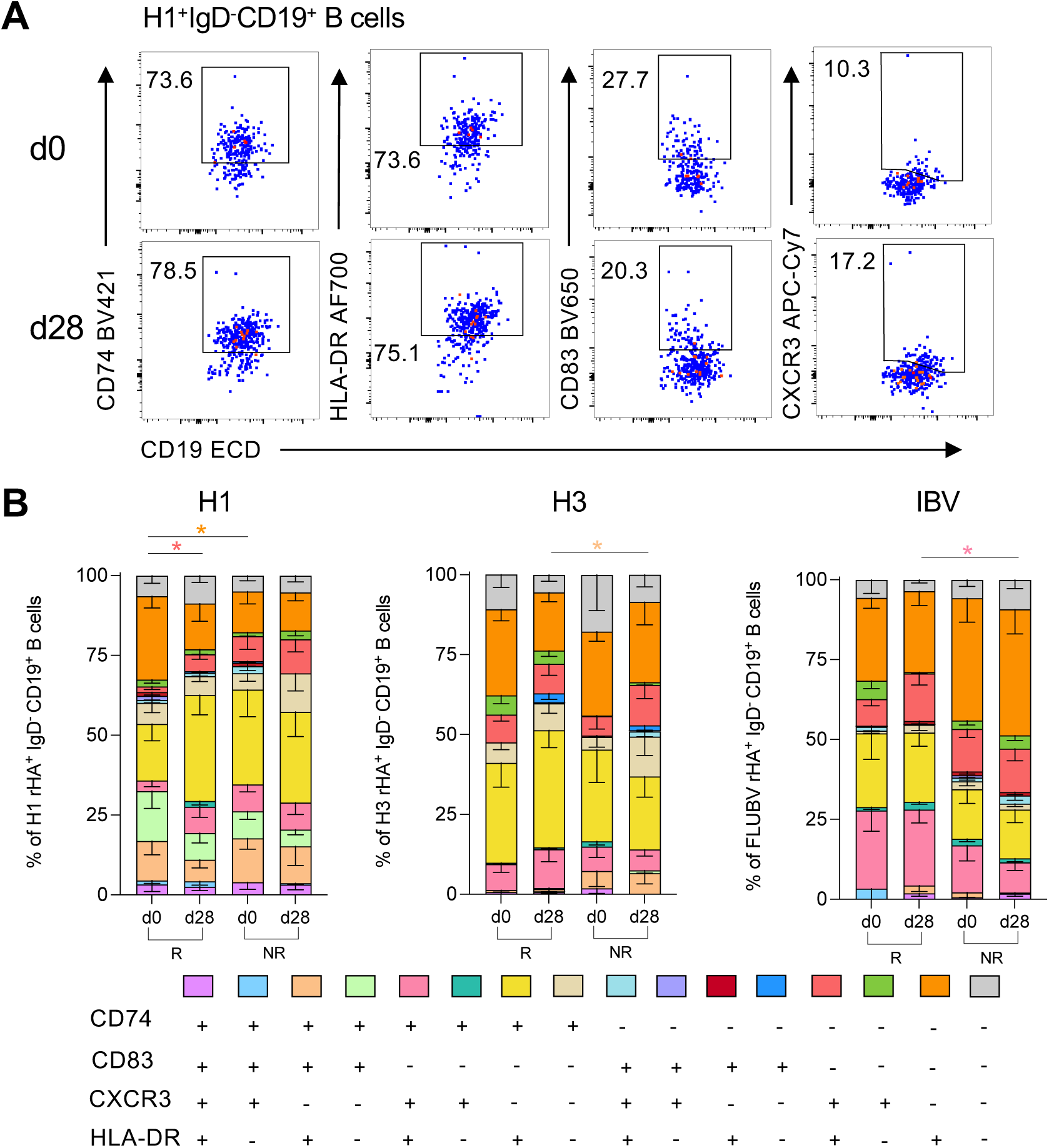
rHA-specific B cells have a unique pattern of CD74, HLA-DR, CD83 and CXCR3 expression. (A) Concatenated FACS plots depicting expression of CD74, HLA-DR, CD83 and CXCR3 for representative H1 specificity (n=19). (B) Co-expression patterns of CD74, HLA-DR, CD83 and CXCR3 on rHA^+^IgD^-^CD19^+^ B cells. Participants in the vaccination cohort with at least 5 and more rHA^+^ B cells were included in the analysis. Statistical significance was assessed using Wilcoxon test for paired timepoint comparisons within groups and Mann-Whitney test for unpaired comparisons between groups (*P<0.05). Mean value plotted, error bars indicate SEM.

Our flow cytometry analyses of CD74, CD83, HLA-DR and CXCR3 expression patterns within influenza rHA+ B cells in Responders and Non-Responders revealed significantly higher frequencies of single-positive HLA-DR^+^ probe-specific H1^+^ B cells at baseline in the H1 Responder group compared to Non-Responders (P=0.0127) (Fig. 8B). Furthermore, within the H1 Responder group, we detected significantly more HLA-DR^+^CXCR3^+^ double-positive B cells on d28 compared to d0 (P=0.0312).

IBV Responders had significantly more HLA-DR^+^CD74^+^CXCR3^+^ rHA+ B cells on d28 compared to Non-Responders (P=0.0179), whereas H3 Responders had significantly less (P=0.0330) HLA-DR^+^CD74^+^CD83^+^ on d28 compared to Non-Responders (Fig. 8B).

Overall, our study demonstrates differential CD74, CD83, HLA-DR and CXCR3 expression patterns in influenza vaccine Responders and Non-Responders, with IIV Responders displaying increased proportions of HLA-DR^+^ H1^+^ B cells at d0 and HLA-DR^+^CXCR3^+^ H1^+^ B cells on d28; increased HLA-DR^+^CD74^+^CXCR3^+^ IBV rHA+ B cells on d28, but decreased HLA-DR^+^CD74^+^CD83^+^ H3^+^ B cells on d28 post-vaccination in comparison to Non-Responders. These data reveal not only B cell phenotypic differences between Responders and Non-Responders but also across vaccine components, thus influenza viral strains.

### Patients hospitalized with influenza have high proportions of atypical B cells and differential expression of CD74, CD83, HLA-DR and CXCR3

Having analyzed CD74, CD83, HLA-DR and CXCR3 expression patterns in IIV immunized participants, we next determined CD74, CD83, HLA-DR and CXCR3 phenotype in patients with acute influenza virus infection in their whole blood samples. We recruited 16 hospitalized influenza patients (10 males and 6 females) in 2025 influenza season in Australia as well as 8 healthy control participants (5 males and 3 females; sex and age-matched to hospitalized participants) (Fig. 9A, B). Hospitalized influenza patients had either acute influenza A (IAV; 11 individuals) or influenza B (IBV; 5 individuals) at the time of sampling (Supplementary Table S8). To define the phenotype of naïve, memory, and atypical B cells in hospitalized influenza patients in comparison to healthy participants, firstly class-switched (IgD^-^CD19^+^) B cells were analysed for expression of CD27 and CD21 (Fig. 9C). We found significantly increased proportion of atypical-like CD27^-^CD21^-^ B cells in hospitalized influenza patients, when compared to healthy participants in IgD^-^ CD19^+^ B cells (healthy mean: 3.7%, infection mean: 8.0%, P=0.0028).

**Figure 9.**
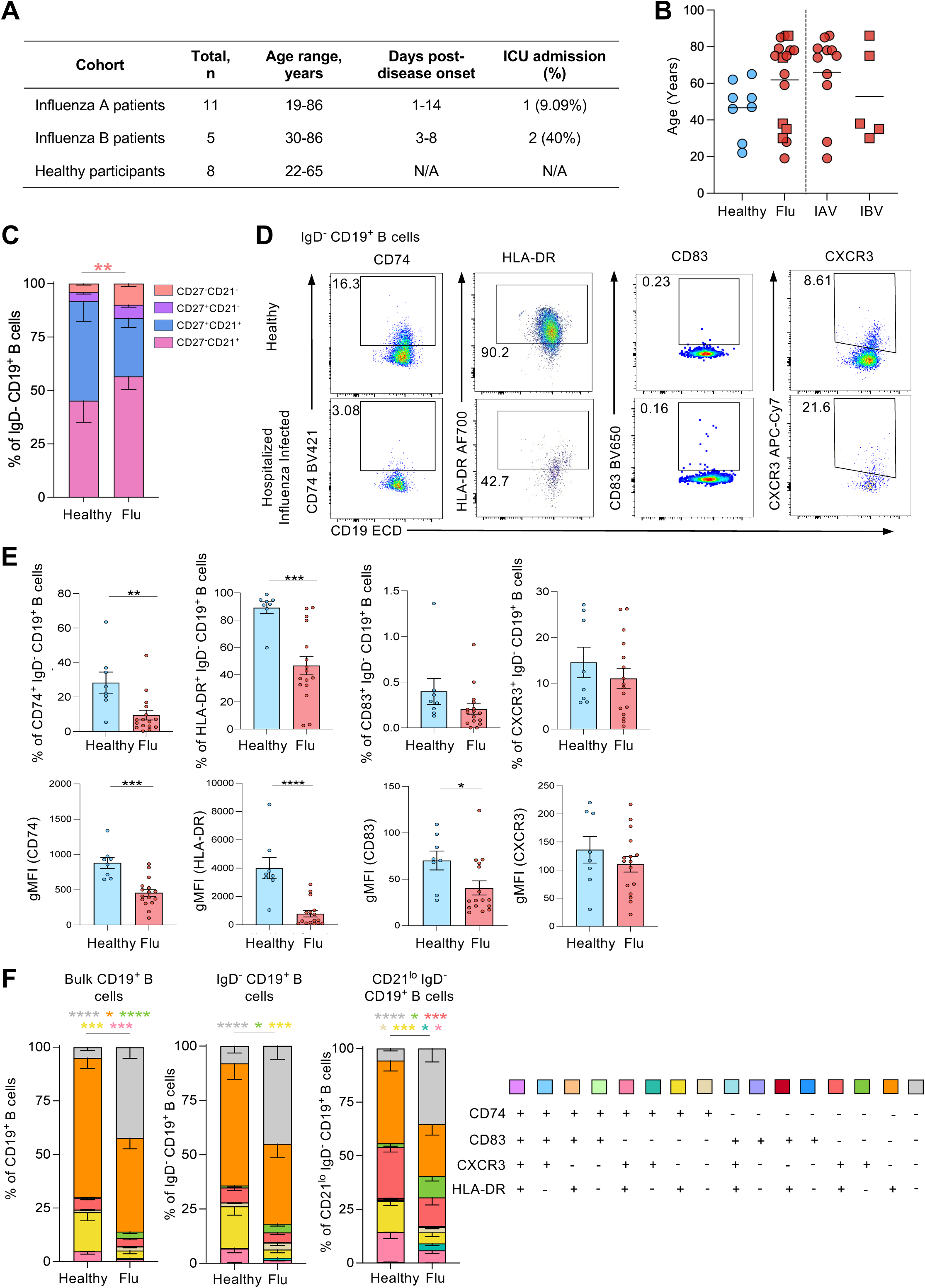
Hospitalized patients with influenza virus infection display higher proportions of atypical B cells than healthy participants. (A) Clinical details of hospitalized influenza patients and healthy controls used in whole blood staining. (B) Age distribution between healthy and infected groups and IAV/IBV distribution in hospitalised patients. (C) Distribution of naïve, memory and atypical B cell phenotypes in class-switched IgD^-^CD19^+^ B cells of healthy and hospitalized influenza virus-infected patients. (D) Representative FACS plots of CD72, HLA-DR, CD83 and CXCR3 expression on IgD^-^CD19^+^ B cells in healthy controls and hospitalized influenza patients. (E) Frequencies CD74+, HLA-DR+, CD83+ and CXCR3+ IgD^-^CD19^+^ B cells and gMFI of CD72, HLA-DR, CD83 and CXCR3 on IgD^-^CD19^+^ B cells in healthy controls and hospitalized influenza patients. Bars indicate mean ± SEM. (F) Co-expression patterns of CD74, HLA-DR, CD83 and CXCR3 in hospitalised influenza patients compared to healthy controls. Boolean gating of these markers was performed on bulk CD19^+^ B cells, IgD^-^CD19^+^ B cells and atypical-like CD21^lo^IgD^-^CD19^+^ B cells. Statistical significance was assessed using Mann-Whitney test (*P<0.05, **P<0.01, ***P<0.001, ****P<0.0001). Error bars indicate SEM.

We examined the expression of CD74, HLA-DR, CD83, and CXCR3 on class-switched IgD^-^CD19^+^ B cells within whole blood of hospitalized influenza patients and healthy participants (Fig. 9D and E). The frequency of CD74^+^ B cells was significantly reduced in hospitalized influenza patients compared to healthy participants (healthy mean: 21.85%, infection mean: 9.51%, P=0.0036), as well as the gMFI of CD74 (healthy mean: 950.1, infection mean: 495.3, P=0.001) (Fig. 9E). This was accompanied by significantly reduced frequency and gMFI of HLA-DR-expressing cells (healthy mean: 89.1% and 4013.3 gMFI; infection mean: 46.6% and 775.9 gMFI; P<0.001). gMFI of CD83 on class-switched B cells was also significantly reduced in hospitalized influenza patients compared to healthy participants (healthy mean: 78.6%, infection mean: 48.5%, P = 0.0324) (Fig. 9E).

We subsequently examined co-expression patterns of CD74, CD83, CXCR3, and HLA-DR on bulk CD19+, IgD^-^CD19^+^ and CD21^lo^IgD^-^CD19^+^ B cells in hospitalized influenza patients compared healthy individuals (Fig. 9F). We found a significantly higher proportion of CD74^-^CD83^-^CXCR3^-^HLA-DR^-^ B cells in the bulk CD19^+^ (P<0.0001), class-switched (IgD^-^CD19^+^) (P<0.0001) and atypical-like (CD21^lo^IgD^-^CD19^+^) B cells (P=0.0002) in hospitalized patients. Additionally, we detected higher proportions of CXCR3^+^ B cells across all three B cell populations (P<0.0001, P=0.0274, P=0.0185), and CXCR3^+^CD74^+^ CD21^lo^IgD^-^CD19^+^ B cells (P=0.0228) in hospitalized influenza patients compared to healthy participants. Conversely, there were significantly lower proportions of HLA-DR^+^ and HLA-DR^+^CD74^+^ B cells across all three B cell populations in influenza patients. We also found significantly lower proportions of CD74^+^CXCR3^+^HLA-DR^+^ bulk CD19^+^ (P=0.0009) and CD21^lo^IgD^-^CD19^+^ B cells (P=0.0192), and CD74^+^ single-positive CD21^lo^IgD^-^CD19^+^ B cells (P=0.0272) in hospitalized influenza patients (Fig. 9F).

Overall, our study demonstrates, for the first time, that hospitalized patients with acute influenza virus infections, for both IAV and IBV, have higher proportions of atypical-like B cells, a finding previously not described for acute influenza infection. Furthermore, we identified changes to the expression patterns of key markers identified in our transcriptomics analyses (CD74, CD83, HLA-DR and CXCR3), potentially affecting B cell proliferation, activation and migration capacity.

## DISCUSSION

Seasonal influenza epidemics cause up to 650,000 deaths annually worldwide {Dhakal, 2019 #5478. Although seasonal influenza vaccines are the best way to combat annual influenza epidemics, the effectivness of influenza vaccines can be low and variable across years and vaccine components, Thus, it is of great importance to understand immune mechanisms driving protective and non-protective responses to IIV. Our scRNA-seq and flow cytometric profiling of rHA-specific and bulk B cells in influenza-vaccinated individuals, vaccinated individuals with confirmed influenza breakthrough infections and hospitalized influenza patients, provides key insights into the immune responses in IIV Non-Responder and Responder vaccinees. Our transcriptomic and phenotypic analyses of influenza HA-specific B cells of IIV Responders and Non-Responders revealed high proportions of atypical A/H1N1- and A/H3N2-specific B cells in Non-Responders compared to Responders at baseline. Hospitalized influenza patients had also higher proportions of atypical B cells compared to healthy participants, a finding not previously reported in acute influenza.

scRNA-seq analysis revealed that both the influenza vaccination cohort and influenza virus breakthrough infection cohort demonstrated distinct subsets of atB cells with a CD21^lo/-^CD27^-^CD11c^+^ phenotype consistent with previous reports {Sutton, 2021 #5454}. We sought to further understand the role of at B cells in responses to IIV, and in acute viral infection. We found significantly higher proportions of A/H1N1- and A/H3N2-specific B cells either single-positive or double-positive for CD11c and FcRL-5 at baseline in Non-Responders compared to Responders, suggesting that the presence of atB cells at baseline may impact the effectiveness of humoral immune responses to IIV. A study investigating humoral responses to the SARS-CoV-2 AZD1222 vaccine with an mRNA vaccine booster found that elderly participants had lower neutralizing antibody responses, associated with higher frequencies of CD27^-^CD21^-^CD11c^+^FcRL-5^+^ atB cells ^38^. A significant negative relationship between atB cell frequency and serum-neutralising activity was found, suggesting that atB cells generate antibodies with lower neutralizing capability, reducing the effectiveness of protective vaccine responses. This could potentially explain why we observed higher frequencies of IAV HA-specific atypical B cells in Non-Responders. During acute influenza, hospitalized patients also had significantly higher proportions of CD27^-^CD21^-^ atB cells.

Research into the role of atB cells in the context of viral infection has thus far been limited to chronic infections. The HIV-1 appears to be a driver of atB cell expansion ^39^ and their production of antibodies with lower neutralization potential ^40^. In malaria, atB cells with high FcRL-5 expression had reduced capacity of secreting antibodies in response to BCR crosslinking compared to FcRL-5-negative atB cells ^41^. It is, however, unknown whether atB cells also increase in frequency during acute viral infections. Our findings of significantly increased CD27^-^CD21^-^ atB cells in hospitalized influenza patients suggest that the atB cells can be elicited by influenza virus infections, closely resembling atB cells described in HIV-1 settings. Further investigation is needed to determine whether atB cells contract following resolution of acute viral infection, and whether disease severity has any effect on the frequency and phenotype of atB cells during acute influenza.

Our scRNA-seq analysis identified four potential markers linked to the atB cells and vaccine responsiveness, namely CD74, HLA-DR, CD83, and CXCR3. CD74 is expressed largely on antigen-presenting cells and is a type II transmembrane protein, which functions as an MHC II chaperone ^42^. Surface CD74 acts as a receptor for members of the macrophage migration inhibitory factor (MIF) cytokine superfamily ^43^. The frequency of CD74-expressing IgD^-^CD19^+^ B cells was significantly reduced in hospitalized influenza patients compared to healthy participants, suggesting that B cells in these patients may have lower rates of proliferation and higher rates of migration ^44,45^. Consistent with lower CD74 expression, the frequency of HLA-DR^+^ B cells was also significantly reduced. MHC II signalling from B cells is required for the maintenance of influenza-specific tissue resident CD4^+^ T cells ^46^, and Epstein-Barr virus downregulates both MHC II and CD74 in CD19^+^ B cells, as a mechanism of evading host immunity ^47^. Furthermore, CD74 is lower in patients with sepsis, associated with poor prognosis ^48^. These findings suggest that CD74 could be a marker of severe outcomes of influenza virus infection, and due to its observed co-expression with MHC II, may have implications for reduced efficacy of the B cell-CD4 T cell axis of immune responses to influenza viruses. We did not find differences in CD74 expression in rHA-specific B cells of vaccinees, suggesting that the role of CD74 may be critical in the outcomes of influenza virus infection rather than vaccination.

We identified CD83 in atypical B cell clusters from the influenza virus breakthrough infection cohort. CD83 is a highly glycosylated type 1 transmembrane glycoprotein part of the immunoglobulin superfamily and is a well-characterized marker for mature and activated dendritic cells ^49^. CD83 is expressed in low levels on B cells after acquiring functional BCRs and plays a role in B cell activation and longevity ^50^, and when blocked, B cells become unresponsive to antigen ^51^. Furthermore, CD83 can be a regulator of B cell functions *in vivo,* as shown in a mouse model of leishmaniasis ^52^. There are limited data regarding CD83’s role in influenza virus infection in humans, although in mice CD83 was shown to regulate antibody production, where CD83 knockout mice had very low IgG titers and delayed B cell recovery in the bone marrow following IAV infection ^53^. Our data show CD83 expression was significantly reduced in hospitalized influenza patients compared to healthy controls, in agreement with the studies in mice ^52^.

CXCR3 can be expressed in atB cells ^54^, potentially as a marker of atB cell activation rather than a lineage-defining gene ^23^. We observed that CXCR3 expression was highly variable between participants in influenza HA-specific and bulk B cells, and the frequency of CXCR3^+^ B cells between resting memory, activated memory, and atypical phenotypes, as well as between timepoints, was comparable.

We observed differential CD74, MHC II, and CXCR3 co-expression patterns in hospitalized influenza patients. Both the bulk and class-switched B cell populations had significantly lower proportions of HLA-DR^+^CD74^+^ double-positive B cells, and significantly higher proportions of CD83^-^CD74^-^CXCR3^-^HLA-DR^-^ quadruple-negative B cells compared to healthy participants. Interestingly, CD74 surface expression can be independent of HLA-DR and HLA-DQ expression in B cells ^55^. Additionally, although there were no differences in overall CXCR3 expression by IgD^-^CD19^+^ B cells in hospitalized influenza patients compared to healthy participants, there were significantly higher proportions of CXCR3^+^ single-positive bulk, class-switched and atypical-like B cells within hospitalized influenza patients than healthy participants. Frequencies of CD74^+^CXCR3^+^ atypical-like B cells were significantly higher in hospitalized influenza patients compared to healthy participants, which when combined with our findings of higher CXCR3 expression on atB and memory B cells, and reports of CXCR3 expression on activated atB and memory B cells ^23,54^, suggests that the atypical-like and memory B cells in hospitalized influenza patients are more activated than those from healthy participants.

Overall, our findings demonstrate increased frequency of atB cells specific for IIV components in Non-Responders, as well as in influenza patients requiring hospitalization. Furthermore, our study suggests that CXCR3 may be a signature of memory and atB cell activation and implicates reduced CD74 and CD83 levels as signatures of perturbed humoral immune responses to influenza virus infection. Collectively, our findings provide new insights underpinning humoral immune responses to IIV and influenza virus infection, and highlight the importance of understanding the role of atypical B cells when evaluating future vaccine design and regimens.

## METHODS

### Study approvals and ethics statement

Influenza vaccination (FLUVAX) and respiratory viral infection cohorts (LIFT, ISSIS, DISI) recruited adults >18 years of age; healthy vaccinees or hospitalized patients, respectively. These studies were approved by the University of Melbourne (31236, 13344, 1443389), Monash Health (HREC/15/MonH/64), Melbourne Health (2016.196), Austin Health (SSA/28204/Austin-2022) and the Alfred Hospital (280/14) Human Research Ethics Committees. In Hong Kong the PIVOT cohorts (ClinicalTrials.gov NCT02957890) recruited community-dwelling older adults aged 69–82 years from the 2017/2018 Northern Hemisphere vaccination season, while the RETAIN cohort (NCT03330132) recruited community-dwelling older adults aged 70–79 years from the 2016/2017 Northern Hemisphere vaccination season. The study protocols were approved by the Institutional Review Board of the University of Hong Kong (PIVOT UW:16-2014, RETAIN UW:16-202) and the University of Melbourne (25684). All participants provided written informed consent prior to inclusion in the study. Experiments conformed to the Declaration of Helsinki Principles and the Australian National Health and Medical Research Council Code of Practice.

### Blood processing

Whole blood was collected in heparinized blood tubes. Plasma was obtained from whole blood in heparinized tubes by collecting supernatant after centrifugation (300x *g*, 10 min, 23°C). PBMCs were isolated through density-gradient centrifugation (Ficoll-Paque, GE Healthcare, Uppsala, Sweden) and cryopreserved in fetal calf serum with 10% DMSO until used in experiments.

### Influenza HA-specific probes

rHA probes were biotinylated tetramerized HA molecules with a removed transmembrane domain and a mutation at Y98F (influenza A virus probes only) to prevent nonspecific binding to sialic acid residues on non-antigen specific B cells ^56^. rHA probes were matched to vaccine components according to the year. H1 (A/Victoria/2570/2019 for 2022, A/Sydney/5/2021 for 2023) and H3 (A/Darwin/6/2021 both years) virus strain rHA probes were each conjugated to a single fluorochrome, whereas the B/Victoria lineage rHA probe (B/Austria/1359417/2021 both years) was conjugated to two fluorochromes, thus B/Victoria rHA^+^ B cells were gated on dual colour positivity.

### Immune profiling flow cytometry analysis using cryopreserved PBMCs

Cryopreserved PBMCs were thawed in cRPMI with Benzonase Nuclease (Merk Millipore, MA, USA) and stained with immunophenotyping antibody staining panels (Supplementary Tables S9 and S10), in some experiments along with rHA probes (Supplementary Tables S11), then fixed in 1% PFA and washed in MACS buffer (PBS with 0.5% BSA and 2 mM EDTA) before samples were acquired on LSR Fortessa (BD Biosciences) and analysed with FlowJo10 (FlowJo, LLC). Gating strategy is provided in Supplementary Figure S1.

### Flow cytometry analysis using whole blood

The antibody mix (Supplementary Table S12) was added directly to 200 µL of whole blood and incubated for 30 min. Red blood cells were lysed with 1X BD FACS Lysing solution (Becton Dickinson, NJ, USA) for 10 min at ambient temperature. Samples were resuspended in 1% PFA for 20 min at 4°C, before washing and resuspending in MACS buffer for acquisition on a BD LSRFortessa II. Gating strategy is provided in Supplementary Figure S2.

### scRNA-seq of PBMCs

PBMCs were thawed in cRPMI with Benzonase Nuclease (Merk Millipore, MA, USA), washed and resuspended in 50µL of MACS buffer. Samples were sequentially stained at 4°C and in the dark with 5µL of TruStain FcX (Biolegend, CA, USA), incubated for 10 min; donor-specific HA probes (Supplementary Table S13) for 15 min, fluorescent monoclonal antibody mix (Supplementary Table S14) for 30 min; CITEseq™-antibodies (Biolegend, CA, USA) (Supplementary Table S15) and TotalSeq anti-human hashtag antibody (where applicable, Biolegend, CA, USA) for 30 min. Samples were washed, resuspended with MACS buffer before sorting into RPMI with 10% fetal calf serum on a FACS Aria III (Becton Dickinson, NJ, USA). Gating strategy for the sort is provided in Supplementary Figure S3. Cells were counted and mixed into 10x Chromium reactions before being loaded into Chromium Next GEM Chip K (PN 2000182, CA, USA), then run on the Chromium Controller iX according to manufacturer’s instructions (Chromium Next GEM Single Cell 5’ v2 (Dual Index) with Feature Barcoding technology; Revision F). Gene expression (GEX), V(D)J dual index and feature barcode (CITE-seq) libraries were generated with Chromium Next GEM Single-Cell 5′ v2 kits according to manufacturer’s instructions. Sequencing was performed using an Illumina NextSeq 500 following manufacturer’s instructions.

### scRNA-seq data analysis

Raw sequencing data were processed with Cell Ranger v7.1.0 (10x Genomics). Individual participants were demultiplexed based on differences in single nucleotide polymorphisms using cellsnp-lite v1.2.2 and vireo v0.2.3 ^57,58^. Gene expression (GEX) and Feature Barcode (CITE-seq) matrices were analyzed with the Seurat package v4.3.0.1 ^59^. Gene expression data were filtered to retain high-quality cells by removing those with mitochondrial percentages or gene counts greater than three Median Absolute Deviations (MADs) above the median, and cells with fewer than 300 detected genes. Sample demultiplexing was performed using the HTODemux function. Each assay was normalized using the default Seurat NormalizeData function. Gene expression data were subsequently scaled using ScaleData applied to variable features only, whereas ADT data were scaled across all features. Principal components were calculated using the 2,000 most variable features following removal of all TCR and BCR genes and the gene MTRNR2L8, which was found to drive unwanted cluster formation. An initial RNA-only UMAP and nearest neighbour graph were constructed using the first 30 principal components for preliminary visualization. Weighted nearest neighbour (WNN) Uniform Manifold Approximation and Projection (UMAP) was subsequently performed to generate final cell clusters by integrating gene expression and protein-level data, incorporating the first 30 and 8 principal components from the RNA and ADT assays, respectively. The FindMarkers function from Seurat was used to identify differentially expressed genes (DEGs) between clusters to help in assigning them to a relevant cell subset. Group analysis was performed using a pseudo-bulk approach after differential expression analysis was performed using the edgeR package ^60^. Data were fitted with a negative binomial generalized log-linear model implemented in glmFit function^61^. Genes were considered differentially expressed between groups if they achieved a FDR <5% after likelihood ratio tests.

### BCR analysis

For BCR data, filtered contigs from the cellranger output were processed using in-house scripts ^62^. Clonal expansions of B cells (>1 clonotype) were calculated using the Immcantation R package (v0.3.2) ^63^. “HierarchicalClones” function of the Scoper sub-package was used for B cell clonal definition by hierarchical clustering with paired heavy and light chain sequences ^63^. Circos plots were plotted using R packages circlize ^64^. Filtered contig outputs from CellRanger (v7.0.0) were read into R and split based on chain type (either IGH or IG[LK]). Contigs were ordered in descending order by UMI count and then collapsed by cell barcode. A full join on cell barcode was performed to obtain paired chain information. The dataset was then filtered by removing cells with missing chain information and cell expressing multiple copies either the light or heavy chains. A threshold of 0.16 was set for eh HierarchicalClones function as well. BCR and GEX data were integrated in Seurat (version 5.2.1) using R (version 4.4.1) with tidyverse (version 2.0.0) packages. BEAM-Ab Assembly probe-binding thresholds were set for each sample using the distribution of probe-specific barcode counts. B cells above threshold for more than one probe, or below threshold for all probes were discarded from repertoire analysis. B cells below threshold for all probes but sharing a clone ID with a probe-binding B cell were retained and assigned to that probe specificity. Circos plots coloured by variable gene usage were generated using circlize version 0.4.16 ^64^ and alluvial plots using ggalluvial version 0.12.5.

### Statistical Analysis

Statistical significance (P<0.05) was assessed using a two-tailed Mann–Whitney U-test, two-tailed Wilcoxon signed-rank test, Fisher’s exact test and two-way ANOVA in GraphPad Prism V10.4.1.

## Acknowledgments

This research was funded in whole or part by the National Health and Medical Research Council Investigator Grants: Investigator L2 to KK (#2033783); L3 to SJK (#2016491), L1 to AKW (#2026762); L1 to THON (#2041012), EL1 to LCR (#2026357) and EL2 to SAV (#2007929). The PIVOT and RETAIN studies were supported by the US Centers for Disease Control and Prevention (Cooperative Agreement Number IP001064-02). We thank all patients and healthy participants who provided blood to our study. We thank all the Research Nurses at Alfred Hospital, Austin Health and Royal Melbourne Hospital who helped with patient recruitment. The authors would like to thank A/Prof Ashraful Haque and Dr Hyun Jae Lee for helpful discussions regarding 10x transcriptomics. We acknowledge the Melbourne Cytometry Platform (The Doherty Institute, UoM) for provision of flow cytometry services. For the purposes of open access, the author has applied a CC BY public copyright licence to any Author Accepted Manuscript version arising from this submission.

## Data availability

All data generated or analyzed during this study are included in this published article (and its supplementary information files) or deposited online. scRNAseq data were deposited at the Gene Expression Omnibus (https://www.ncbi.nlm.nih.gov/geo/) with accession number: GSE310898.

## Notes

### Competing Interest Statement

Katherine Kedzierska has received paid honoraria from Pfizer. Hayley McQuilten has a consultancy role for Ena Therapeutics

