## Supplementary tables-1 for "Defining influenza-specific B cells in vaccine responders, non-responders and influenza breakthrough infections"

**Supplementary Table S5**. Common genes identified in the Responder groups and shared between different specificities.

| **Groups** | **Number of common elements** | **Common elements** |
| --- | --- | --- |
| H3N2, H1N1, B/Vic and B/Yam | 0 |  |
| H3N2, H1N1 and B/Yam | 0 |  |
| H3N2, B/Vic and B/Yam | 0 |  |
| H1N1, B/Vic and B/Yam | 4 | *AK8, LCK, SH2D3A, PRIM1* |
| H3N2, H1N1 and B/Vic | 10 | *CNTNAP2, SETD9, NEIL1, XXYLT1, AC253572.2, IFNG-AS1, TSGA10, FGD6, EPHA4, ABHD5* |
| H3N2 and B/Yam | 2 | *MTRNR2L8, TXNL4B* |
| H1N1 and B/Yam | 1 | *TSPAN13* |
| H3N2 and B/Vic | 20 | *NUDT19, LINS1, S100Z, NBPF12, TMEM191B, METRN, MFAP1, POC1B-AS1, GDPD5, NLK, MOB1B, TEX9, ARNT, AC245014.3, ADCK2, ZSWIM9, SBF2, AL136419.3, RAD17, CAMK4* |
| H1N1 and B/Vic | 19 | *HLA-DQB2, SGPP2, AC092944.1, ZNF528, BRIP1, AC006115.2, SPATA24, AC135507.1, TMEM53, MARC2, HTR3A, DNAJC5B, NIF3L1, BTN3A2, SCCPDH, THAP7-AS1, KLF8, PRMT7, DSE* |
| H3N2 and H1N1 | 22 | *PTPDC1, CAMK2N1, FCHO2, AL022332.1, AL356512.1, AL049830.3, FAM50B, AL034397.3, ZNF862, AC104596.1, PIK3R4, TCF7, CCDC127, LINC00324, ZNF322, KCNA3, CRIP2, ASB16-AS1, CYP2U1, GPR160, AC100810.1, BEND5* |
| B/Vic and B/Yam | 19 | *HLA-DQA2, PTOV1-AS1, CPM, GPR155, AHI1, LINC00996, BRF2, GK, SLC38A11, DHFR, FLVCR1-DT, AC124248.1, BTNL9, SYNPO, CYP4V2, TNFRSF17, PASK, EID2, UTP14A* |

**Supplementary Table S6**. Common genes identified in the Non-Responder groups and shared between different specificities.

| **Groups** | **Number of common elements** | **Common elements** |
| --- | --- | --- |
| H3N2, H1N1, B/Vic and B/Yam | 0 |  |
| H3N2, H1N1 and B/Yam | 0 |  |
| H3N2, B/Vic and B/Yam | 7 | *TOP1MT, ATP13A3, SERTAD3, NR4A1, DUSP5, AREG, TUBB2A* |
| H1N1, B/Vic and B/Yam | 1 | *RTP5* |
| H3N2, H1N1 and B/Vic | 19 | *TIMP1, CLEC2B, METRNL, LINC01128, KANSL1-AS1, CEMIP2, DGKG, RGS2, NR4A2, SERTAD1, TRIB1, AC104024.1, MT1X, GRAP2, PTPN7, SEMA4A, EGR1, CMTM3, MCOLN1* |
| H3N2 and B/Yam | 0 |  |
| H1N1 and B/Yam | 0 |  |
| H3N2 and B/Vic | 17 | *HAUS3, PLOD3, PANK3, AC021678.2, UBALD1, TRPS1, AC058791.1, GIT1, IQGAP2, KCNQ1OT1, IGHM, EPG5, DUSP2, WASF1, TYW1B, AC233755.1, ZNF551* |
| H1N1 and B/Vic | 12 | *MOXD1, MPP6, HSPB1, ARSD, ENC1, IQCN, AC026356.2, CREB3L2, POLR2J2, SPON2, NR4A3, ACE* |
| H3N2 and H1N1 | 35 | *ZEB2, CD72, TENT5A, ARHGAP18, CBLB, BATF, CLN8, HCST, SNTB2, PLD4, CAMK2D, MAP4K3-DT, SOX5, ZNF749, ZBTB32, PLEKHG2, RGS1, MIR4435-2HG, SLC11A1, MT2A, PCDH9, LILRB2, CDK5R1, PEBP4, MAP4K3, ITGAX, TCF19, BCL11B, CPQ, ARL4D, MAP2, SERPINB8, GPR137B, TAS1R3, CHI3L2, TRPS1, AC058791.1, GIT1, IQGAP2, KCNQ1OT1, IGHM, EPG5, DUSP2, WASF1, TYW1B, AC233755.1, ZNF551* |
| B/Vic and B/Yam | 19 | *CD83, LRRC40, KIAA0930, JUNB, C1RL-AS1, MAPK6, BEND4, PER1, PTPRO, HHLA3, SH3RF1, AC240274.1, NEXN, SOCS3, PLTP, LCMT2, PPP1R16A, RETREG1, AC008074.2* |

**Supplementary Table S7**. Fluvax participants used in scRNAseq and flow cytometry experiments and their vaccine responder status.

|  | **A/H1N1** | **A/H3N2** | **B/VICTORIA** | **B/YAMAGATA** |
| --- | --- | --- | --- | --- |
| **Participant ID** | A/Victoria/2570/2019 | A/Darwin/9/2021 | B/Austria/1359417/2021 | B/Phuket/3073/2013 |
| Participant 1 | L-R | H-R | L-R | H-R |
| Participant 2 | L-R | L-NR | H-NR | H-NR |
| Participant 3 | L-R | L-R | L-R | H-NR |
| Participant 4 | L-NR | L-NR | H-NR | H-NR |
| Participant 5 | L-R | L-NR | L-R | L-R |
| Participant 6 | H-R | H-R | L-R | H-NR |
| Participant 7 | H-NR | L-NR | L-R | H-NR |
| Participant 8 | L-R | L-R | H-R | H-R |
| Participant 9 | H-NR | L-R | L-R | H-R |
| Participant 10 | H-R | L-R | L-R | H-NR |
| Participant 11 | H-NR | L-R | L-NR | H-NR |
| Participant 12 | H-NR | L-R | L-NR | H-NR |
| Participant 13 | H-NR | L-R | L-NR | H-NR |
| Participant 14 | L-R | L-R | L-NR | H-NR |
| Participant 15 | H-NR | L-R | L-R | H-R |
| Participant 16 | L-R | H-R | L-NR | H-NR |
| Participant 17 | H-NR | L-NR | H-NR | H-NR |
| Participant 18 | H-NR | H-NR | H-NR | H-NR |
| Participant 19 | H-R | L-R | L-R | L-R |

**Supplementary Table S8**. Hospitalized influenza patients and the healthy controls used in the flow cytometry experiments.

| **Donor ID** | **Age** | **Number of days post-symptom onset at blood collection** | **Diagnosis** | **Ward** |
| --- | --- | --- | --- | --- |
| Flu-1 | 79 | 3 | Influenza A | General |
| Flu-2 | 65 | 3 | Influenza A | General |
| Flu-3 | 86 | 7 | Influenza A | General |
| Flu-4 | 85 | 4 | Influenza A | General |
| Flu-5 | 59 | 6 | Influenza A | General |
| Flu-6 | 19 | 8 | Influenza A | General |
| Flu-7 | 38 | 8 | Influenza B | ICU |
| Flu-8 | 28 | 1 | Influenza A | General |
| Flu-9 | 78 | 14 | Influenza A | ICU |
| Flu-10 | 78 | 4 | Influenza A | General |
| Flu-11 | 75 | 6 | Influenza A | General |
| Flu-12 | 35 | 5 | Influenza B | General |
| Flu-13 | 30 | 6 | Influenza B | General |
| Flu-14 | 86 | 6 | Influenza B | ICU |
| Flu-15 | 74 | 2 | Influenza A | General |
| Flu-16 | 75 | 3 | Influenza B | General |
| Healthy-1 | 52 | N/A | N/A | N/A |
| Healthy-2 | 27 | N/A | N/A | N/A |
| Healthy-3 | 46 | N/A | N/A | N/A |
| Healthy-4 | 62 | N/A | N/A | N/A |
| Healthy-5 | 47 | N/A | N/A | N/A |
| Healthy-6 | 52 | N/A | N/A | N/A |
| Healthy-7 | 65 | N/A | N/A | N/A |
| Healthy-8 | 22 | N/A | N/A | N/A |

**Supplementary Table S9**. Antibody panel for phenotyping influenza HA-specific B cells within PBMCs of participants receiving IIV.

| **Antibody** | **Clone** | **Dilution** | **Company** | **Catalog** |
| --- | --- | --- | --- | --- |
| CD11c FITC | Bu15 | 1:50 | Biolegend | 337214 |
| FcRL-5 RB705 | 509F8 | 1:150 | BD | 757499 |
| IBV rHAs APC | N/A | N/A | N/A | N/A |
| HLA-DR AF700 | G46-6 | 1:50 | BD | 560743 |
| CXCR3 APC-Cy7 | G025H7 | 1:100 | Biolegend | 353722 |
| CD74 BV421 | LN2 | 1:200 | BD | 743731 |
| Streptavidin BV510 | N/A | 1:600 | BD | 563261 |
| LIVE/DEAD Aqua | N/A | 1:500 | Thermofisher | L34966 |
| CD3 BV510 | OKT3 | 1:600 | Biolegend | 317332 |
| CD8 BV510 | RPA-T8 | 1:1500 | Biolegend | 301047 |
| CD10 BV510 | HI10a | 1:750 | Biolegend | 312219 |
| CD14 BV510 | M5E2 | 1:300 | Biolegend | 301841 |
| CD16 BV510 | 3G8 | 1:500 | Biolegend | 302047 |
| CD27 BV605 | O323 | 1:150 | Biolegend | 302830 |
| CD83 BV650 | HB15e | 1:400 | BD | 740602 |
| H1 rHA BV711 | N/A | N/A | N/A | N/A |
| IgG BV786 | G18-145 | 1:75 | BD | 564230 |
| H3 rHA PE | N/A | N/A | N/A | N/A |
| CD19 ECD | J3-119 | 1:100 | Beckman | IM2708U |
| IgD PE-Cy7 | IA6-2 | 1:500 | BD | 561314 |
| IgM BUV395 | G20-127 | 1:150 | BD | 563903 |
| CD21 BUV737 | B-ly4 | 1:300 | BD | 612788 |

**Supplementary Table S10**. Antibody panel for phenotyping B cells within PBMCs of the RETAIN participants.

| **Antibody** | **Clone** | **Dilution** | **Company** | **Catalog** |
| --- | --- | --- | --- | --- |
| CD11c FITC | Bu15 | 1:50 | Biolegend | 337214 |
| FcRL-5 RB705 | 509F8 | 1:150 | BD | 757499 |
| CXCR3 APC-Cy7 | G025H7 | 1:100 | Biolegend | 353722 |
| CD74 BV421 | LN2 | 1:200 | BD | 743731 |
| Streptavidin BV510 | N/A | 1:600 | BD | 563261 |
| LIVE/DEAD Aqua | N/A | 1:500 | Thermofisher | L34966 |
| CD3 BV510 | OKT3 | 1:600 | Biolegend | 317332 |
| CD8 BV510 | RPA-T8 | 1:1500 | Biolegend | 301047 |
| CD10 BV510 | HI10a | 1:750 | Biolegend | 312219 |
| CD14 BV510 | M5E2 | 1:300 | Biolegend | 301841 |
| CD16 BV510 | 3G8 | 1:500 | Biolegend | 302047 |
| CD27 BV605 | O323 | 1:150 | Biolegend | 302830 |
| CD83 BV650 | HB15e | 1:400 | BD | 740602 |
| IgG BV786 | G18-145 | 1:75 | BD | 564230 |
| CD19 ECD | J3-119 | 1:100 | Beckman | IM2708U |
| IgD PE-Cy7 | IA6-2 | 1:500 | BD | 561314 |
| IgM BUV395 | G20-127 | 1:150 | BD | 563903 |
| CD21 BUV737 | B-ly4 | 1:300 | BD | 612788 |

**Supplementary Table S11.** Influenza rHA probes used in phenotyping experiments.

| **Panel** | **Subtype** | **Strain** | **Fluorochrome** |
| --- | --- | --- | --- |
| FLUVAX | H1N1 | A/Sydney/5/2021(H1N1)pdm09-like | BV711 |
| RETAIN | H1N1 | A/California/7/2009 | BV711 |
| RETAIN | H1N1 | A/Michigan/45/2015 | BV711 |
| FLUVAX | H3N2 | A/Darwin/9/2021 | PE |
| RETAIN | H3N2 | A/Switzerland/9715292/2013 | PE |
| FLUVAX | B/Vic | B/Austria/1359417/2021 | APC |
| RETAIN | B/Vic | B/Brisbane/60/2008 | APC |
| FLUVAX & RETAIN | B/Yam | B/Phuket/3073/2013 | APC |

**Supplementary Table S12**. Antibody panel for phenotyping whole blood B cells in patients hospitalized with respiratory viral diseases.

| **Antibody** | **Clone** | **Dilution** | **Company** | **Catalog** |
| --- | --- | --- | --- | --- |
| CD11c FITC | Bu15 | 1:50 | Biolegend | 337214 |
| FcRL-5 RB705 | 509F8 | 1:150 | BD | 757499 |
| HLA-DR AF700 | G46-6 | 1:50 | BD | 560743 |
| CXCR3 APC-Cy7 | G025H7 | 1:100 | Biolegend | 353722 |
| CD74 BV421 | LN2 | 1:200 | BD | 743731 |
| Streptavidin BV510 | N/A | 1:600 | BD | 563261 |
| CD3 BV510 | OKT3 | 1:600 | Biolegend | 317332 |
| CD8 BV510 | RPA-T8 | 1:1500 | Biolegend | 301047 |
| CD10 BV510 | HI10a | 1:750 | Biolegend | 312219 |
| CD14 BV510 | M5E2 | 1:300 | Biolegend | 301841 |
| CD16 BV510 | 3G8 | 1:500 | Biolegend | 302047 |
| CD27 BV605 | O323 | 1:150 | Biolegend | 302830 |
| CD83 BV650 | HB15e | 1:400 | BD | 740602 |
| IgG BV786 | G18-145 | 1:75 | BD | 564230 |
| CD19 ECD | J3-119 | 1:100 | Beckman | IM2708U |
| IgD PE-Cy7 | IA6-2 | 1:500 | BD | 561314 |
| IgM BUV395 | G20-127 | 1:150 | BD | 563903 |
| CD21 BUV737 | B-ly4 | 1:300 | BD | 612788 |

**Supplementary Table S13**. Influenza rHA probes used in 10X scRNA-seq experiment.

| **Virus** | **DNA barcode** | **Catalogue** |
| --- | --- | --- |
| B/Austria/1359417/2021- like strain | TotalSeq™-C0956 APC Streptavidin | 405283 |
| B/Brisbane/60/2008-like virus | TotalSeq™-C0956 APC Streptavidin | 405283 |
| B/Phuket/3073/2013 - like strain | TotalSeq™-C0957 APC Streptavidin | 405285 |
| A/Darwin/9/2021 (H3N2) - like strain | TotalSeq™-C0958 APC Streptavidin | 405293 |
| A/2018/2019 H1 Bris/2018 | TotalSeq™-C0958 APC Streptavidin | 405293 |
| A/California/7/2009 (H1N1)-like virus | TotalSeq™-C0959 APC Streptavidin | 405159 |
| A/Sydney/5/2021 (H1N1)pdm09 – like strain | TotalSeq™-C0959 APC Streptavidin | 405159 |
| B/Brisbane/60/2008-like virus | TotalSeq™-C0960 APC Streptavidin | 405157 |

**Supplementary Table S14**. Antibody panel for FACS sorting of PBMC samples used in 10x scRNA-seq experiment.

| **Antibody** | **Clone** | **Dilution** | **Company** | **Catalogue** |
| --- | --- | --- | --- | --- |
| CD4-BUV496 | SK3 | 1:100 | BD | 612936 |
| CXCR5-BV421 | RF8B2 | 1:25 | BD | 562747 |
| CD71-BV650 | CY1G4 | 1:50 | Biolegend | 334116 |
| CD19-BV711 | SJ25C1 | 1:100 | BD | 563038 |
| CD20-AF700 | 2H7 | 1:100 | BD | 560631 |
| CD8-PerCP Cy5.5 | SK1 | 1:100 | BD | 565310 |
| CD3-PE-CF594 | UCHT1 | 1:100 | BD | 562280 |
| IgD-PE Cy7 | IA6-2 | 1:500 | BD | 561314 |
| CD14-BV510 | M5E2 | 1:100 | Biolegend | 301841 |
| SAV-BV510 |  | 1:600 | Biolegend | 405233 |

**Supplementary Table S15**. CITEseq antibody panel used in 10x scRNA-seq experiment.

| **Antibody** | **Specificity** | **Catalogue** |
| --- | --- | --- |
| TotalSeq™-C0171 | ICOS | 313553 |
| TotalSeq™-C0088 | PD-1 | 329963 |
| TotalSeq™-C0140 | CXCR3 | 353747 |
| TotalSeq™-C0144 | CXCR5 | 356939 |
| TotalSeq™-C0181 | CD21 | 354923 |
| TotalSeq™-C0154 | CD27 | 302853 |
| TotalSeq™-C0046 | CD8 | 344753 |
| TotalSeq™-C0072 | CD4 | 300567 |
| TotalSeq™-C0053 | CD11c | 371521 |
