## Supplementary figures for "Defining influenza-specific B cells in vaccine responders, non-responders and influenza breakthrough infections"

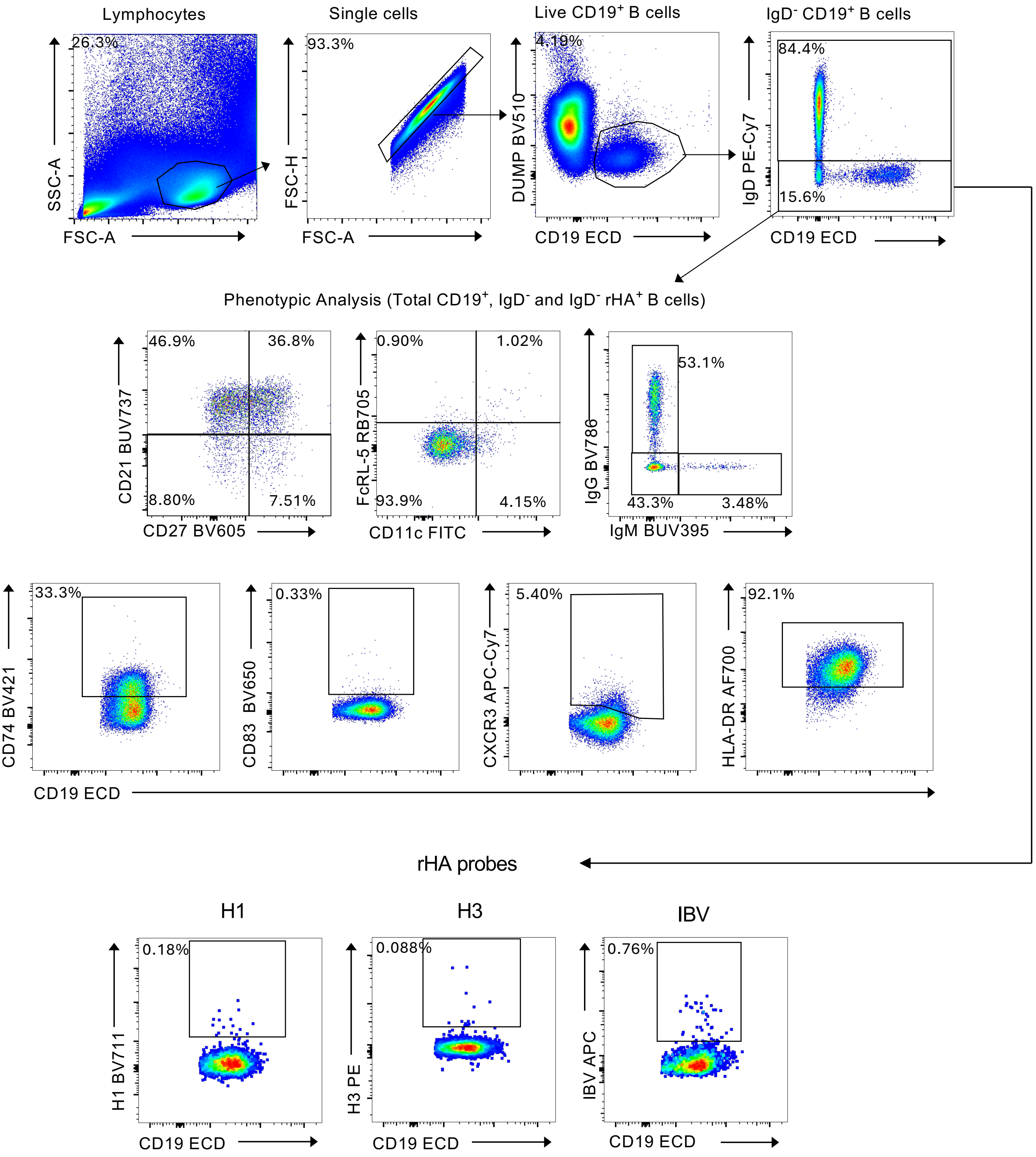


**Supplementary Figure S1** Gating strategy for B cell subsets and influenza HA-specific B cells from Australian influenza vaccinee cohort. B cells were identified as live, CD19$^+$ CD3$^-$ CD8$^-$ CD10$^-$ CD14$^-$ CD16$^-$ single cell lymphocytes not bound to free streptavidin. Class-switched B cells (IgG$^+$, IgM$^+$) B cells were identified within live IgD$^-$ B cells. B cell phenotypes were assessed by expression of CD21 and CD27, CD11c and FcRL-5, CD74, CD83, CXCR3 and HLA-DR. Influenza HA-specific (rHA probe-specific) B cells were identified within live IgD$^-$ B cells. B cell phenotypes were additionally assessed within Influenza HA-specific B cells.

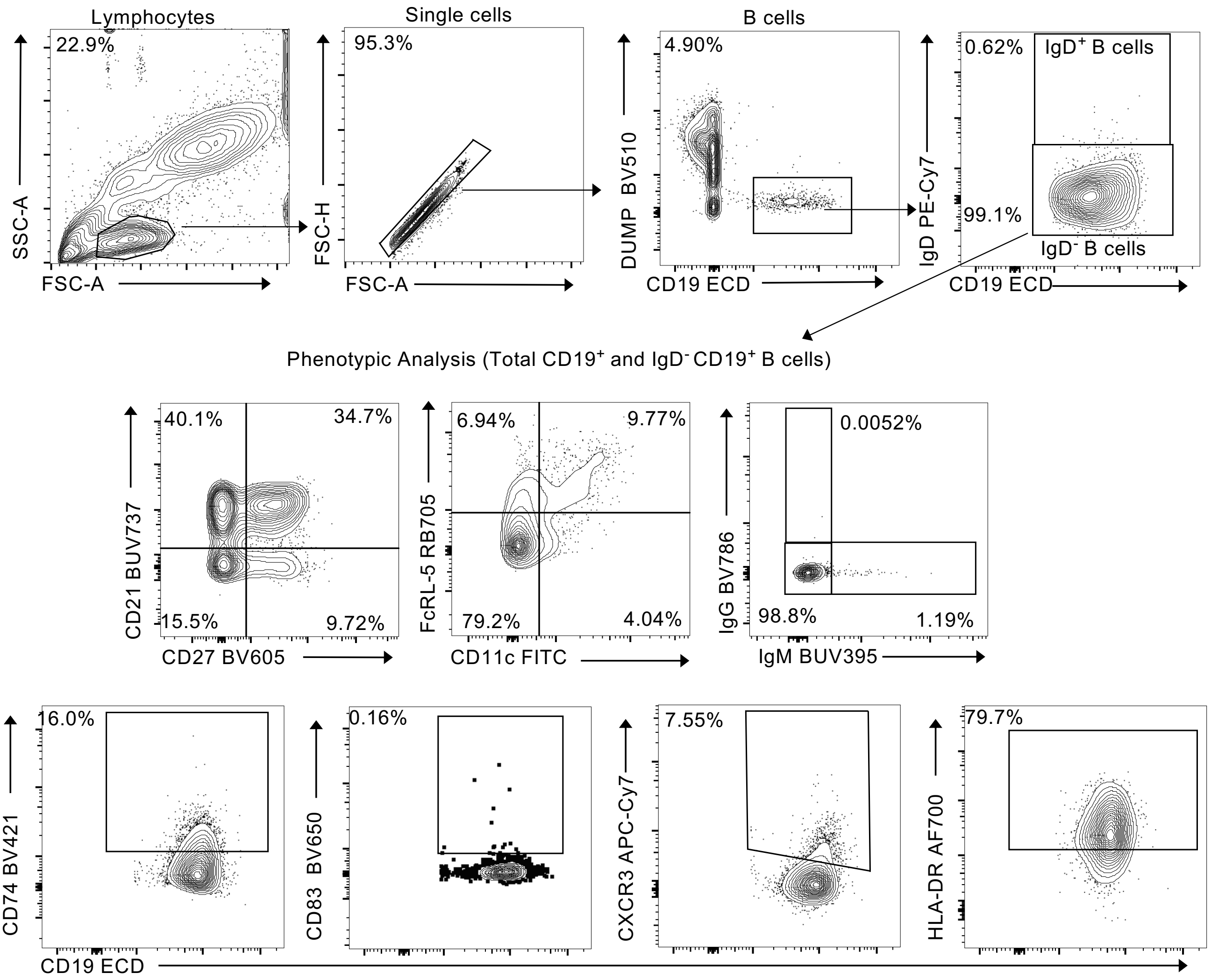


**Supplementary Figure S2** Gating strategy for B cell subsets from whole blood of Australian hospitalized influenza virus infection patients and healthy participants. B cells were identified CD19$^{+}$ CD3$^{-}$ CD8$^{-}$ CD10$^{-}$ CD14$^{-}$ CD16$^{-}$ single cell lymphocytes. Class-switched B cells (IgG$^{+}$, IgM$^{+}$) B cells were identified within IgD$^{-}$ B cells. B cell phenotypes were assessed by expression of CD21 and CD27, CD11c and FcRL-5, CD74, CD83, CXCR3 and HLA-DR.

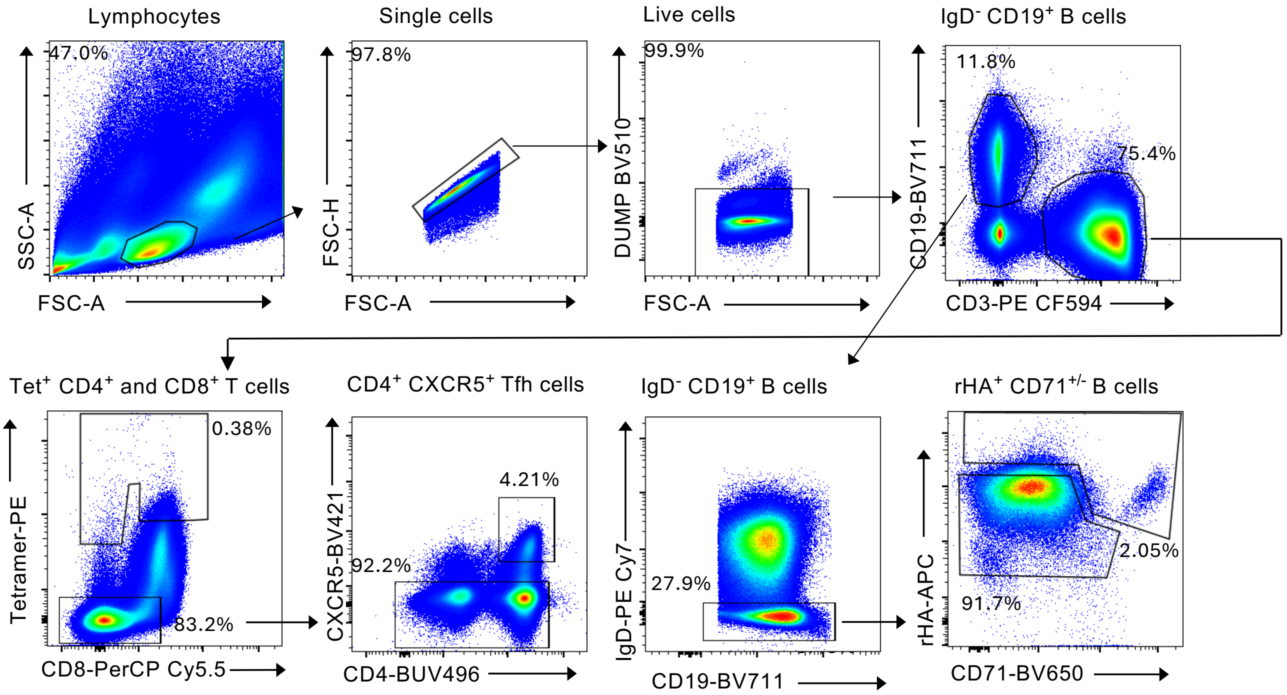


**Supplementary Figure S3** Gating strategy for 10X geonomics sorting of influenza HA-specific B cells, tetramer-positive T cells and CD4$^{+}$CXCR5$^{+}$ Tfh cells from Australian influenza and Hong Kong vaccinee cohorts. B cells were identified as live, CD19$^{+}$CD3$^{-}$CD14$^{-}$ single cell lymphocytes not bound to free streptavidin. Influenza HA-specific (rHA probe-specific) B cells were identified within live IgD$^{-}$ B cells as rHA$^{+}$CD71$^{hi/lo}$. B cell phenotypes were additionally assessed within Influenza HA-specific B cells. Tetramer-positive (both class I and II) T cells were identified as CD3$^{+}$CD19$^{-}$CD8$^{+/-}$ T cells. Tfh cells were identified within live CD8- population as CXCR5$^{+}$CD4$^{+}$ T cells.
